# Imputing abundance of over 2500 surface proteins from single-cell transcriptomes with context-agnostic zero-shot deep ensembles

**DOI:** 10.1101/2024.07.31.605432

**Authors:** Ruoqiao Chen, Jiayu Zhou, Bin Chen

## Abstract

Cell surface proteins serve as primary drug targets and cell identity markers. The emergence of techniques like CITE-seq has enabled simultaneous quantification of surface protein abundance and transcript expression for multimodal data analysis within individual cells. The published data have been utilized to train machine learning models for predicting surface protein abundance based solely from transcript expression. However, the small scale of proteins predicted and the poor generalization ability for these computational approaches across diverse contexts, such as different tissues or disease states, impede their widespread adoption. Here we propose SPIDER (surface protein prediction using deep ensembles from single-cell RNA-seq), a context-agnostic zero-shot deep ensemble model, which enables the large-scale prediction of cell surface protein abundance and generalizes better to various contexts. Comprehensive benchmarking shows that SPIDER outperforms other state-of-the-art methods. Using the predicted surface abundance of >2500 proteins from single-cell transcriptomes, we demonstrate the broad applications of SPIDER including cell type annotation, biomarker/target identification, and cell-cell interaction analysis in hepatocellular carcinoma and colorectal cancer.

## Introduction

Cell surface proteins are crucial for a cell to sense extracellular signals, mediate cell-cell interactions, and perform cellular functions. They often serve as cell identity markers and hold the most significant therapeutic implications, covering over 60% of current drug targets (Santos, Rita, et al., 2017; Bausch-Fluck Damaris et al., 2018). Quantifying surface protein abundance at cellular levels provides broad applications including cell type annotation, disease biomarker discovery, drug target identification, and cell-cell interaction analysis. There has been rapid development in single-cell multi-omics technologies for profiling cell surface protein abundance, including CITE-seq (Stoeckius Marlon et al., 2017), REAP-seq (Peterson Vanessa M et al., 2017), ABseq (Shahi Payam et al., 2017), and mass cytometry-based multimodal assay (Bennett, Hayley M., et al., 2023). These methods use antibodies to tag cell surface proteins and allow for the simultaneous measurement of cell surface protein abundance and transcriptome in the same cell, yet several significant limitations remain. These experiments remain cost-prohibitive in many labs, and technology barriers restrict widespread access to these methods. Moreover, while in theory these technologies have no upper bound for measurable surface proteins, in practice they only routinely measure fewer than 300 surface proteins, and even merely 10∼20 surface proteins in many cases, only accounting for a small portion of the maximum number of 5570 human cell surface proteins as predicted by the Human Protein Atlas (HPA) (Uhlén, Mathias, et al., 2015). This is not solely due to cost considerations but also because many proteins lack suitable antibodies (Vistain, Luke F., and Savaş Tay., 2021).

To mitigate these challenges, computational approaches offer promising options by predicting surface protein abundance from single-cells transcriptomes. Current approaches include multimodal data integration models such as Seurat (Hao, Yuhan, et al., 2021), totalVI (Gayoso, Adam, et al., 2021) and sciPENN (Lakkis, Justin, et al., 2022), as well as models specifically designed for surface protein imputation like cTP-net (Zhou, Zilu, et al., 2020). These models are trained on CITE-seq datasets to learn the relation between transcript expression and surface protein abundance. Yet, the downstream application of their predicted surface protein abundance to addressing real biomedical problems is limited by two unsolved challenges. Firstly, these models are constrained to predicting the same proteins as present in the training/reference CITE-seq dataset, thus limiting the scale of predictable surface proteins to fewer than 300. Secondly, these models do not intend to generalize to various contexts (e.g., different tissues or disease states). In fact, there are much fewer CITE-seq datasets compared to single-cell RNA-sequencing (scRNA-seq) datasets, and CITE-seq datasets for certain tissues and diseases are rare or absent. For instance, there is a lack of CITE-seq dataset for the human kidney or Alzheimer’s disease, making it difficult to prepare a reference dataset that shares the same cellular context as a provided query dataset, highlighting the need for designing models to extrapolate surface protein abundance prediction to different contexts.

Here we propose SPIDER, a context-agnostic zero-shot deep ensemble model that enables abundance prediction for a large scale of cell surface proteins across various contexts with strong generalization ability. To consider the influence of contextual variations on surface protein expression, and improve model performance in new contexts, SPIDER incorporates contextual information, encompassing tissue, disease state, and cell type, as integral components of the input alongside transcriptome data. Moreover, to expand the scale of predictable proteins, SPIDER predicts for both proteins seen during training and unseen proteins (different from the trained proteins) via zero-shot learning mechanisms. Through comprehensive benchmarking, we show that SPIDER outperforms baselines in terms of both seen and unseen proteins in a variety of contexts. Further, we use SPIDER to predict the abundance for over 2500 cell surface proteins on hepatocellular carcinoma (HCC) and colorectal cancer (CRC) liver metastasis transcriptomes. Based on its prediction, we perform downstream analyses including cell type annotation, disease biomarker identification, and cell-cell interaction analysis. These applications underscore SPIDER’s versatility across a wide array of contexts, even those different from the reference dataset.

## Results

### SPIDER model overview

SPIDER employs a deep ensemble architecture combined with zero-shot learning methods to achieve abundance prediction for thousands of surface proteins in single cells (Fig. 1a). To start with, a reference dataset is built from CITE-seq dataset(s) and trained for SPIDER to learn a relation between each specific protein’s cell surface abundance and the single-cell transcriptomes (Fig. 1b). However, such relation could be largely affected by other environmental *context factors* including tissue, disease and cell type, especially considering the situation where a query dataset may source from a context different from the reference dataset, increasing the difficulty for accurate prediction. Therefore, SPIDER also adopts a context-agnostic approach to elevate context generalization ability, where one-hot encoding is used to encode the information on these context factors and combines it with transcriptomes for training (Fig. 1b). To bolster SPIDER’s context-awareness, a comprehensive reference dataset is compiled by consolidating CITE-seq datasets from six studies, which comprise a total of 289 proteins and 120,461 cells covering five tissues, four diseases, and 17 cell types (Methods) (Fig. 2a and Extended Data Table 1). To mitigate batch effects among transcriptome datasets, scArches-SCANVI (Lotfollahi, Mohammad, et al) is used to embed them into lower dimensions before combining with context factors (Fig. 2b, Methods). During SIPDER’s training stage, deep neural networks (DNN) are individually trained for each protein with an internal 10-fold random holdout validation, where each DNN captures the intricate relation between each protein’s cell surface abundance and the embedded transcriptomes within its corresponding context. These trained proteins are referred to as seen proteins, and the trained weights of corresponding DNNs are saved and directly used to predict cell surface abundance of the same 289 proteins on any given embedded query scRNA-seq dataset within its context (Fig. 1b).

**Fig. 1.**
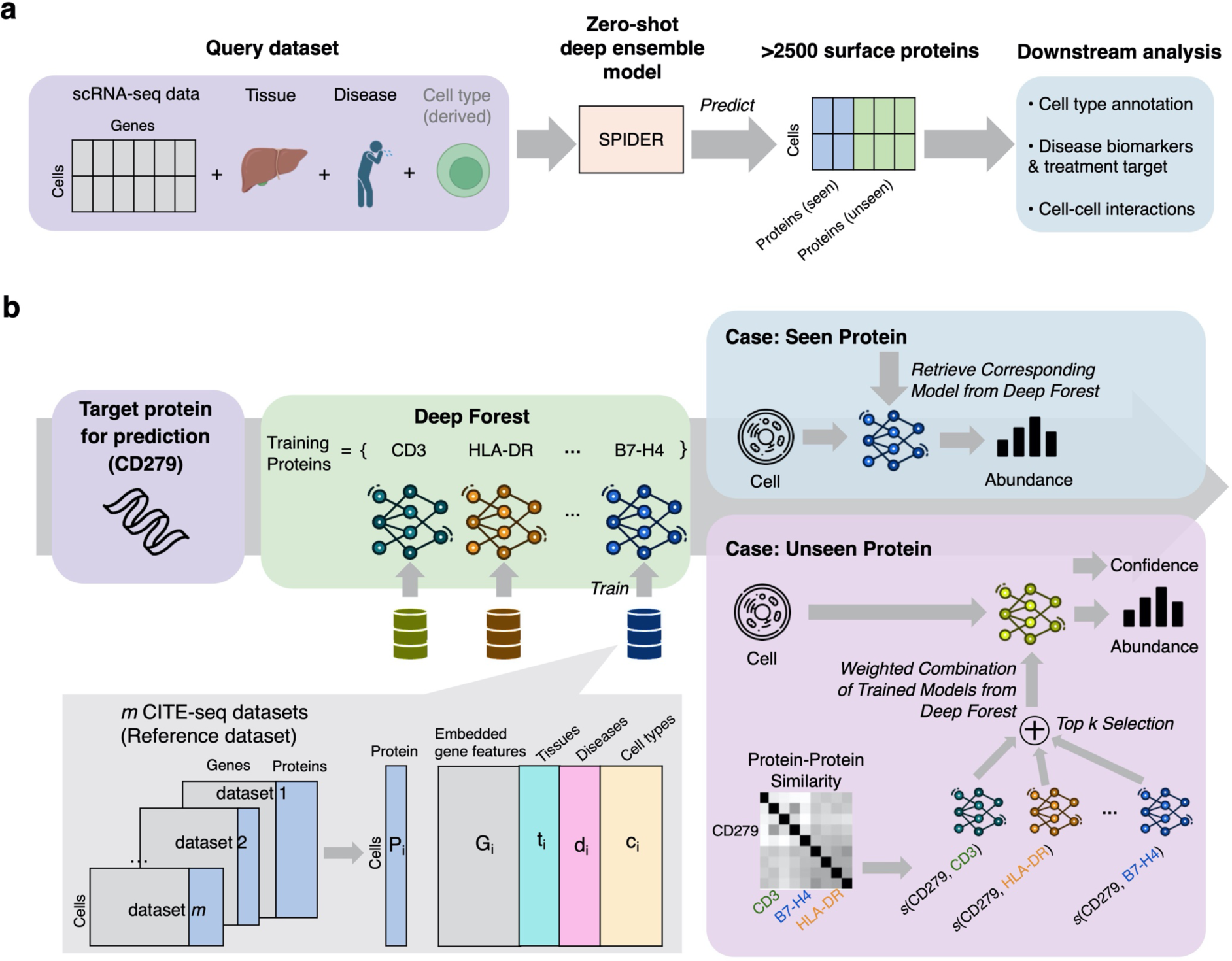
SPIDER analysis pipeline. **a**, SPIDER takes query single-cell transcriptome data and contextual information including tissue, disease and cell type as input, then uses its zero-shot deep ensemble architecture to predict the abundance for large-scale (>2500) cell surface proteins on the query transcriptome data. SPIDER predicts both seen and unseen proteins. SPIDER- predicted surface protein abundance data allow downstream analysis including cell type annotation, disease biomarker identification, and cell-cell interaction analysis. **b**, Schematic of the SPIDER model. SPIDER builds a reference set using m CITE-seq datasets. SPIDER splits the m datasets (Gi, Pi) and combines them with one-hot encoded contextual information (ti, di, ci) to train protein-specific DNNs, which are used to directly predict the abundance for seen cell surface proteins on query transcriptome data. The trained DNNs also form ensemble members for predicting unseen proteins. The deep ensembles are combined with a zero-shot learning method where context-specific protein-protein similarity (S) generated from the query transcriptome data is used as auxiliary information for relating unseen proteins to seen proteins. Finally, SPIDER estimates unseen proteins with maximum protein-protein similarity of >0.85 to be highly confident predicted proteins.

**Fig. 2.**
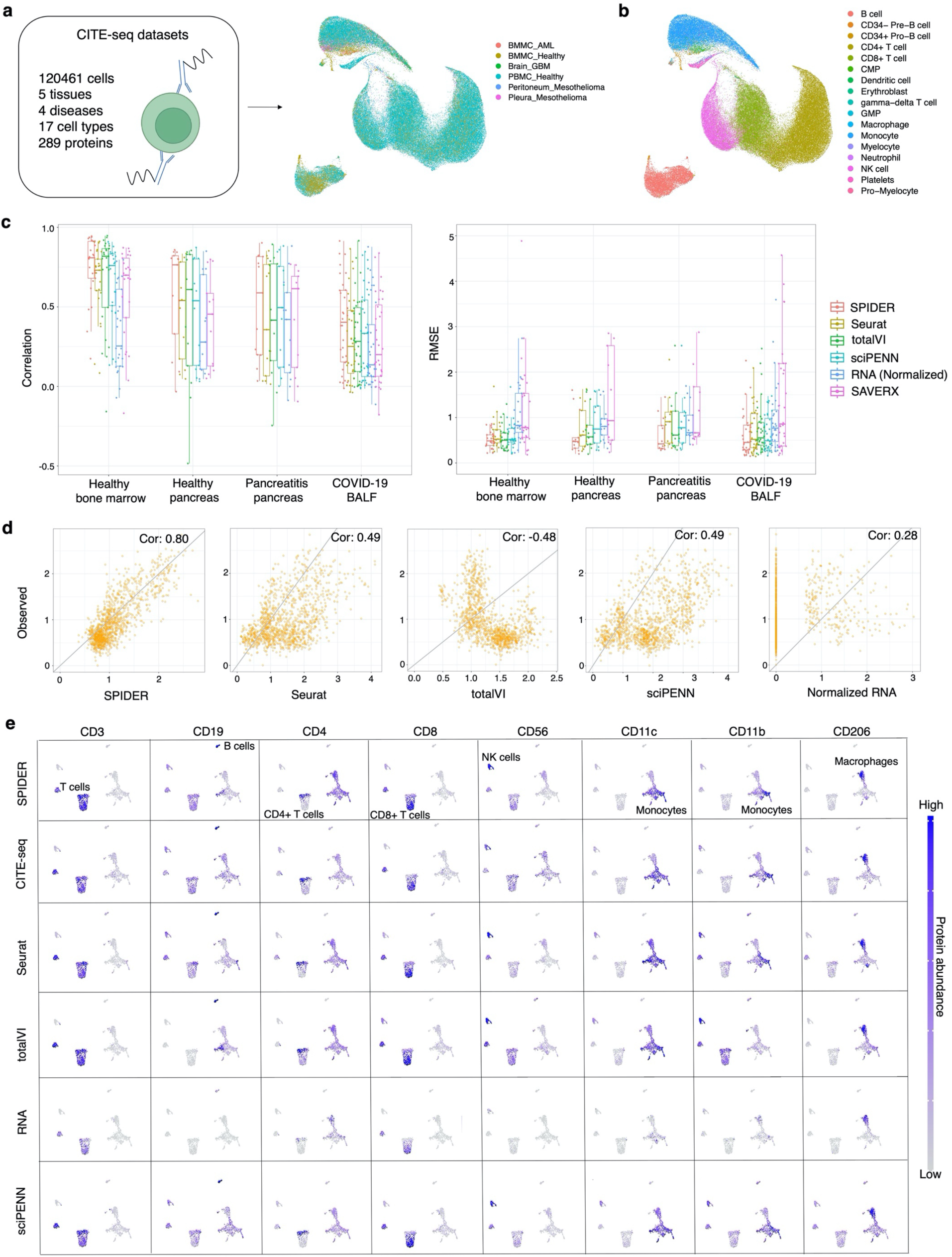
Benchmarking with seen proteins across various contexts. **a-b,** UMAP visualization of the embeddings on the reference set’s transcriptomes. **a,** CITE-seq datasets from various contexts are combined together to construct the reference set. Embeddings of the transcriptomes are visualized and colored by dataset source. **b**, Embeddings of the transcriptomes are visualized and colored by cell type. **c,** Prediction accuracy for every seen protein in all cells. Four CITE-seq datasets covering a variety of contexts are used as external validation datasets: healthy bone marrow, healthy pancreas, pancreatitis pancreas and COVID-19 BALF. SPIDER is compared to Seurat V4, totalVI, sciPENN, normalized RNA expression, and SAVERX-imputed transcriptomes. Pearson correlation (Left) and RMSE (Right) are shown between every cell surface protein’s predicted abundance (or normalized RNA expression) and CITE-seq measured abundance for each method. Each dot represents a protein. **d**, Scatter plots of CD11b’s predicted abundance (or normalized RNA expression) versus CITE-seq measured abundance across all cells in the healthy pancreas query dataset. Each dot represents a cell. **e**, UMAP visualization of the transcriptome data of the healthy pancreas query dataset, colored by predicted abundance (or normalized RNA expression) and CITE-seq measured abundance of selected cell type markers across all cells. All protein abundances are in the log-scale.

To achieve the prediction of the abundance of unseen cell surface proteins, SPIDER enables the zero-shot capability. Zero-shot learning (ZSL) originates from the problem setting where one needs to predict the class for a sample whose actual class was not seen during the training stage (Xian, Yongqin, et al., 2018). ZSL methods generally solve this problem by relating seen and unseen classes through auxiliary information such as class-class similarity. SPIDER extends the ZSL method to solve the regression problem of predicting the abundance for unseen proteins by adopting the protein-protein similarity as a type of auxiliary information that relates a given unseen protein to every seen protein in the reference set (Fig. 1b). To derive context-specific protein-protein similarity, we first gather the names for all seen and unseen proteins, and for each protein we compute its gene co-expression pattern based on the query transcriptome, where the co-expression value is calculated between this protein’s corresponding coding gene and all the other genes in the query transcriptome data (Methods). This gene co-expression matrix then serves as protein representation, and the cosine similarity is calculated between a given unseen protein’s and every seen protein’s representation. After obtaining this protein-protein similarity,

SPIDER then proceeds to perform zero-shot prediction for unseen proteins via a deep ensemble architecture which combines previously trained DNNs with a linear regression algorithm (Fig. 1b). For each given unseen protein, a filtering is applied to the DNNs by first removing the low-quality DNNs with poor internal validation performance, and then ranking the rest of the high-quality DNNs by their corresponding protein-protein similarity to the given unseen protein, where the top eight DNNs with the largest similarity values are selected to form ensemble members for that unseen protein’s prediction (Methods). Each ensemble member makes a separate prediction on the query dataset, and the outputs have their weights assigned the same values as their corresponding protein-protein similarities. The outputs from ensemble DNNs are aggregated using these weights to produce the final prediction of the given unseen protein’s cell surface abundance on the query dataset (Fig. 1b).

In addition, SPIDER also estimates the prediction confidence for all the seen and unseen proteins, where higher confidence means the protein is more likely to be predicted accurately (Fig. 1b). A seen protein is estimated to be highly confident if its corresponding DNN’s internal 10-fold random holdout validation performance shows a correlation exceeding 0.6 between prediction and ground truth, and an unseen protein is estimated to be highly confident if it has a maximum protein-protein similarity of >0.85 to the ensemble members (Methods).

### Prediction performance for seen proteins across various contexts

In order to elevate model generalization ability across various contexts, besides transcriptomes, we also incorporate three types of *context factors*: tissue, disease and cell type (Fig. 1b). To validate that adding *context factors* to the input indeed improves model performance, we compare the internal validation performance of SPIDER for seen proteins with different model input combinations. Results show that SPIDER performs best when all three types of *context factors* are added to the input (Extended Data Fig. 1). Only adding tissue or disease factor to model input also elevates model performance compared to utilizing transcriptomes alone. Interestingly, the inclusion of cell type factor does not yield a discernible impact on model performance within this internal validation, likely because the expression features embedded from scArches-SCANVI have already encoded pertinent cell type information (Extended Data Fig. 1).

To assess SPIDER’s capability in predicting the abundance of a large scale of surface proteins across datasets with various contexts, including those differing from the reference dataset, we conduct external validation on four CITE-seq datasets covering a variety of contexts: healthy bone marrow, healthy pancreas, pancreatitis pancreas, and COVID-19 bronchoalveolar lavage fluid (BALF), respectively (Ordered from the most similar to the reference dataset, to the least similar) (Extended Data Table 1). Since SPIDER utilizes different prediction approaches for seen and unseen proteins, and the baselines for comparison are also different, we evaluate the prediction accuracy for seen proteins and unseen proteins separately.

We first consider the condition of predicting for seen proteins on the four external validation sets. To assess whether SPIDER’s performance is satisfying or not, we compare SPIDER’s prediction accuracy to state-of-the-art surface protein prediction methods Seurat V4, totalVI, sciPENN and cTPnet (Fig. 2c and Extended Data Fig. 2). Finally, we add another two baselines, one using each protein’s corresponding normalized RNA count as the prediction, the other one using imputed RNA values by SAVERX (Wang, Jingshu, et al., 2019), as when scRNA-seq experiments are conducted in lieu of CITE-seq experiments, normalized or imputed transcriptomes are often used by researchers as a surrogate for representing protein abundance.

We evaluate the prediction performance for seen proteins from several different perspectives. Firstly, we evaluate the prediction performance for every seen protein across all cells. For all four tested external validation sets, SPIDER achieves higher median correlation and lower median RMSE between the prediction and ground truth of the tested seen proteins, compared to the Seurat, sciPENN, cTPnet and normalized RNA baselines (Fig. 2c and Extended Data Fig. 2). For comparison to totalVI and SAVERX baselines, SPIDER also outperforms them in terms of both median correlation and RMSE for three out of all four external validation sets, only obtaining a slightly lower median correlation in one dataset, where SPIDER still outperforms them in terms of median RMSE (Fig. 2c). Notably, there is a tendency that the more different the query dataset’s context is from the reference set, the larger degree SPIDER outperforms other baselines. As shown in Fig. 2c, compared to the second best-performed model totalVI, SPIDER elevates the correlation by 25.5%, 39.8%, 42.5%, for the healthy pancreas, pancreatitis pancreas, and COVID-19 BALF query datasets, respectively. This indicates that SPIDER’s context-agnostic approach largely improves the model’s ability to generalize across diverse contexts compared to other models. The scatter plot of CD11b’s prediction in all cells on the query healthy pancreas dataset illustrates an example of the significant improvement in prediction accuracy by SPIDER compared to other baselines, where SPIDER’s prediction is highly correlated to the ground truth (correlation coefficient = 0.80), while none of the three baselines reach a correlation coefficient of 0.5 (Fig. 2d).

Secondly, we evaluate the within-cell type prediction performance for every seen protein. Cell types containing at least 100 cells are selected for evaluation. For the COVID-19 BALF query dataset, in all three cell types, SPIDER outperforms Seurat, totalVI and normalized RNA in terms of both median correlation and median RMSE (Extended Data Fig. 3a). For the other three query datasets, SPIDER consistently exhibits better overall performance in most cell types compared to the other baselines (Extended Data Fig. 3b-d).

Thirdly, we examine the distribution of predicted abundance for several cell type markers selected from the seen proteins across all cells. In the healthy pancreas query dataset, visualization of SPIDER’s predicted surface protein abundance across all cells closely resembles that of CITE-seq (Fig. 2e). Compared to normalized RNA, all four machine learning models show stronger signals of expression at the similar magnitude as CITE-seq. Moreover, visualization of the aforementioned protein CD11b shows evident high abundance only in monocytes for CITE-seq quantification and SPIDER prediction, whereas Seurat, totalVI and sciPENN falsely predict high abundance of CD11b in NK cells and T cells as well, indicating that SPIDER captures cell marker abundance distributions more accurately than the other baselines (Fig. 2e).

Taken together, these results prove that SPIDER is capable of predicting cell surface abundance for seen proteins in query datasets with similar or even completely different contexts from the reference dataset, and that SPIDER achieves higher prediction accuracy than current state-of-the-art methods. Another interesting observation is that in three out of all four query datasets, compared to SPIDER, totalVI displays a wide range of correlations among all predicted seen proteins, with its lowest correlation significantly lower than that of SPIDER (Fig. 2c). This suggests that SPIDER is more robust for prediction than totalVI.

### SPIDER enables accurate prediction for unseen proteins

One strength of SPIDER is its capability of predicting cell surface abundance for a large scale of proteins. This achievement goes beyond the prediction for seen proteins and encompasses the prediction of considerable unseen proteins. Protein-protein similarity serve as auxiliary information in our ZSL method for predicting unseen proteins (Fig. 1b). Considering the inherent variability in expression patterns of cell surface proteins across different query dataset contexts, SPIDER uses an across-all-cell type gene co-expression pattern generated from the query transcriptomes as protein representations to further derive context-specific protein-protein similarity (Methods). During the process of determining the best type of protein representations, we also consider using Gene Ontology (GO) terms, Protein-Protein Interaction (PPI) scores, as well as within-cell type gene co-expression patterns generated from the query transcriptomes as another three types of protein representations to further derive the protein-protein similarity, where the first two approaches barely associate with the query dataset’s context (Methods). Our comparison results of internal validation performance for unseen proteins show that SPIDER performs the best when using the across-all-cell type context-specific gene co-expression pattern as protein representations, with the median correlation between prediction and ground truth elevated by 158.9%, 84.7%, and 75.80% compared to using GO terms, PPI scores, and within-cell type gene co-expression patterns, respectively (Extended Data Fig. 4).

To evaluate SPIDER’s prediction accuracy for unseen proteins in various contexts, we utilize the same four external validation sets as used in the evaluation of seen proteins. For each protein in an external validation set that has its corresponding RNA expression measured, we first deliberately make it an unseen protein by excluding its corresponding DNN from the prediction process entirely, and then select the ensemble members solely from the remaining trained DNNs for this unseen protein’s prediction. We repeat this set-aside and prediction process to predict for every protein in all four external validation sets and further compare SPIDER’s prediction accuracy to the RNA baseline. We exclude Seurat, totalVI and cTPnet from baselines since they are incapable of predicting unseen proteins.

For unseen proteins we also evaluate the prediction performance from aforementioned three aspects as with seen proteins. Firstly, by evaluating the prediction performance for every protein in all cells, our results show that in all four query datasets, among all the tested unseen proteins, SPIDER achieves a higher median correlation between prediction and ground truth compared to the RNA baseline (Fig. 3a). It could be expected that the overall prediction accuracy of unseen proteins should be poorer than that of seen proteins, as no ground truth labels of unseen proteins appear during training. Nevertheless, SPIDER manages to reach a moderately strong median correlation of 0.59, 0.56 and 0.53 for the healthy bone marrow, healthy pancreas and pancreatitis pancreas datasets, respectively (Fig. 3a). Regarding median RMSE, SPIDER also outperforms the RNA baseline in three out of all four datasets, and performs comparably to the RNA baseline for the healthy bone marrow dataset (Fig. 3a). By selecting highly confident unseen proteins estimated by SPIDER (Methods), we could enhance the median correlation to 0.81, 0.63, 0.68, and 0.79 with a protein coverage of 41.8%, 58.3%, 50.0% and 6.7% for the respective query datasets (Fig. 3b). For the selected highly confident proteins, SPIDER continues to surpass the RNA baseline in three out of all four query datasets regarding the correlation metric, and in all four query datasets regarding the RMSE metric (Fig. 3b). Furthermore, in two out of all four query datasets, all the selected highly confident proteins could achieve a relatively accurate prediction with a correlation coefficient of >0.5 between predicted abundance and ground truth (Fig. 3b).

**Fig. 3.**
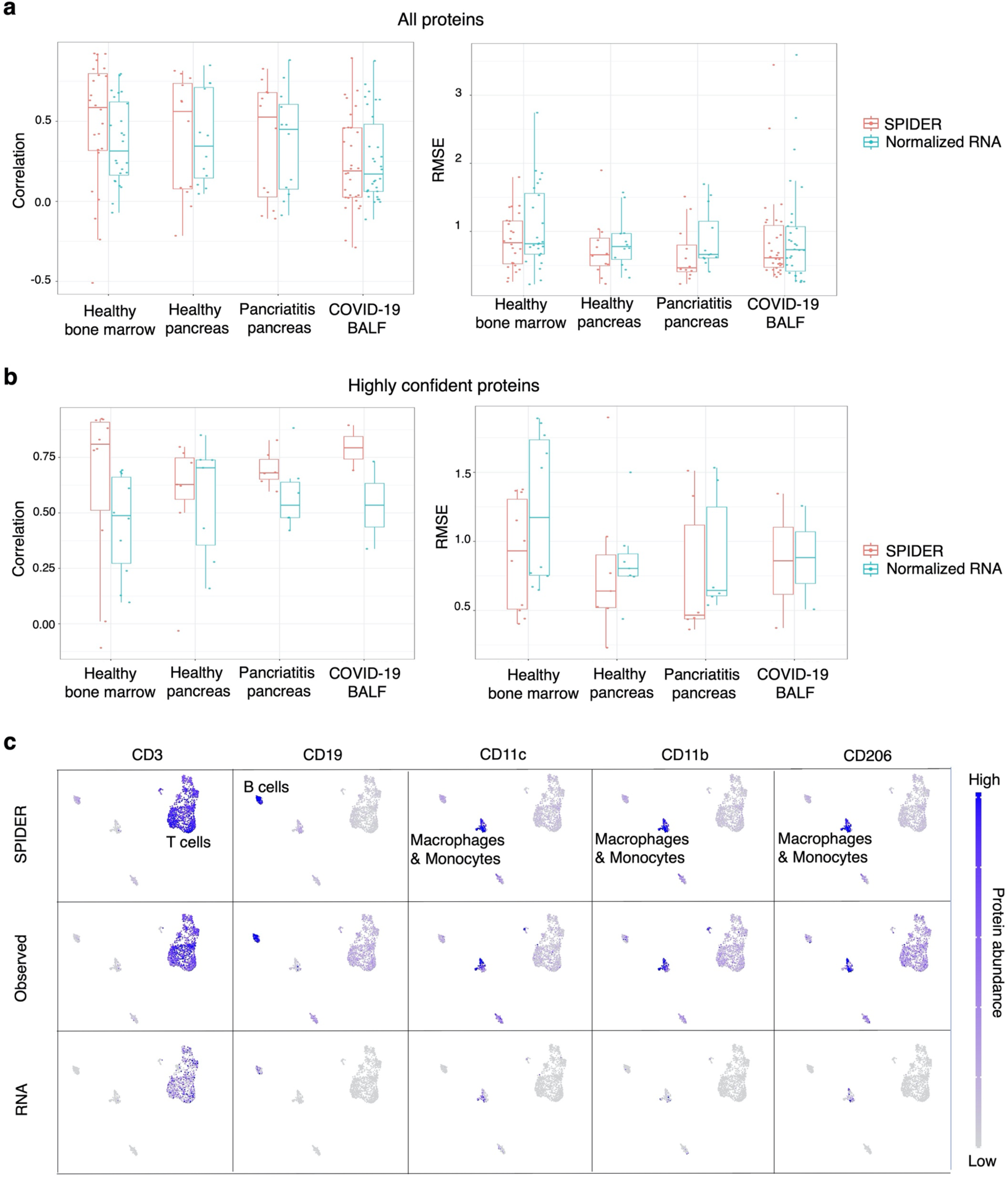
Benchmarking with unseen proteins across various contexts. **a,** Prediction accuracy for every unseen protein in all cells. The same four CITE-seq datasets as with the evaluation of the seen proteins are used as external validation datasets. SPIDER is compared to normalized RNA expression. Pearson correlation (Left) and RMSE (Right) are shown between every cell surface protein’s predicted abundance (or normalized RNA expression) and CITE-seq measured abundance for each method. Each dot represents a protein. **b**, Prediction accuracy for every SPIDER-estimated highly confident unseen protein across all cells. SPIDER is compared to normalized RNA expression. Pearson correlation (Left) and RMSE (Right) are shown between every cell surface protein’s predicted abundance (or normalized RNA expression) and CITE-seq measured abundance for each method. Each dot represents a protein. **c**, UMAP visualization of the transcriptome data of the pancreatitis pancreas query dataset, colored by predicted abundance (or normalized RNA expression) and CITE-seq measured abundance of selected cell type markers across all cells. All protein abundances are in the log-scale.

Secondly, we evaluate the within-cell type prediction performance for every tested unseen protein. In the healthy bone marrow query dataset, SPIDER outperforms the RNA baseline in 11 out of all 12 cell types in terms of median RMSE, and in 7 out of 12 cell types in terms of correlation (Extended Data Fig. 5a). In the other three query datasets, SPIDER performs better than the RNA baseline regarding RMSE in most of the cell type (Extended Data Fig. 5b-d).

Thirdly, we scrutinize the distribution of predicted abundance for several cell type markers selected from SPIDER’s estimated highly confident unseen proteins. Results show that in the pancreatitis pancreas query dataset, visualization of SPIDER’s predicted surface protein abundance across all cells closely resembles that of CITE-seq (Fig. 3c). Compared to RNA, SPIDER shows stronger signals of expression, exhibiting the similar magnitude as CITE-seq (Fig. 3c). Taken together, our results demonstrate that beyond seen proteins, SPIDER is equipped to predict abundance for unseen proteins while ensuring high accuracy through confidence estimation.

### SPIDER assists with cell type annotation

To showcase SPIDER’s various downstream applications, from scRNA-seq data we first apply SPIDER to predict the abundance for all the human cell surface proteins as listed by the UniProt database, and then conduct multiple downstream analysis using the predicted surface protein abundance values (Methods). We first aim to demonstrate SPIDER’s application to cell type annotation, an important procedure for characterizing cells and understanding cellular heterogeneity within cell populations. We use two pre-integrated and annotated scRNA-seq datasets sourcing from human liver with distinct disease contexts of (1) hepatocellular carcinoma and (2) healthy, respectively (Fig. 4d-e) (Lu, Yiming, et al., 2022) (Annotation is performed in a manuscript under submission). We utilize SPIDER to predict the abundance for 2811 surface proteins on these two datasets (Extended Data Table 1, Supplementary Table 1).

**Fig. 4.**
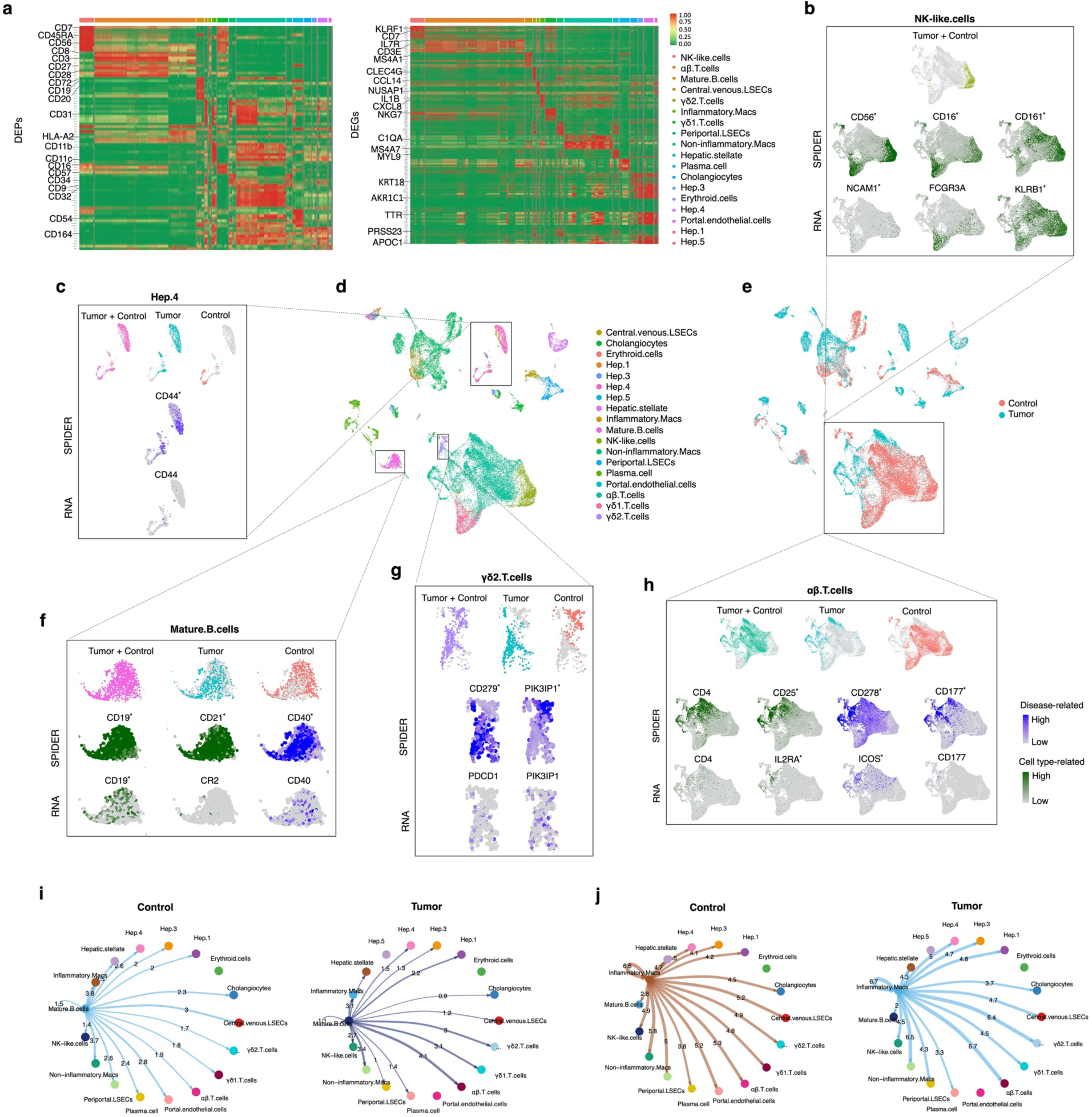
Application of SPIDER to human hepatocellular carcinoma scRNA-seq data. **a,** Heatmaps showing the top DEPs (Left) or DEGs (Right) with maximum fold-change identified using differential expression analysis for each cell type on SPIDER-predicted surface protein abundance and transcriptome data, respectively. **b-h**, comparison between SPIDER-predicted surface protein abundance and corresponding RNA expression level for identified cell type-related or disease-related proteins within specific cell populations. Disease-related surface proteins and their corresponding RNA are colored dark-green, and cell type-related surface proteins are colored dark-purple. Surface proteins and genes labeled with asterisk mean they are DEPs or DEGs under corresponding differential expression analysis. **b**, UMAP visualization of predicted surface protein abundance and corresponding RNA expression in NK-like cells. **c**, UMAP visualization of predicted surface protein abundance and corresponding RNA expression in Hep.4. **d-e**, UMAP visualization of all cells in the HCC dataset, colored (**d**) by cell type or (**e**) by disease state. Hep: Hepatocytes. Macs: Macrophages. LSECs: Liver sinusoidal endothelial cells. **f-h**, UMAP visualization of predicted surface protein abundance and corresponding RNA expression (**f**) in mature B cells, (**g**) in γδ2 T cells, and (**h**) in αβ T cells. **i-j**, Cell-cell interactions inferred from SPIDER-predicted surface protein abundance. The values represent the strength of interactions. **i**, CCIs between mature B cells and other cell populations under tumor and control conditions. **j**, CCIs between inflammatory macrophages and other cell populations under tumor and control conditions.

To demonstrate SPIDER’s application to cell type annotation, one prerequisite is to confirm that SPIDER can accurately identify the canonical surface markers for corresponding cell types, as these markers are commonly used to define a cell type for each cell cluster during annotation. Thus, we conduct differential expression (DE) analysis among the pre-annotated cell types based on predicted protein abundance levels or RNA levels, leading to differentially expressed proteins (DEPs) or differentially expressed genes (DEGs) serving as markers for these distinct cell types (Methods, Supplementary Table 1). Our results show that many canonical cell type surface markers could be identified by SPIDER, for instance, CD56, CD16, CD161, CD7 for NK-like cells, CD3, CD8, CD27, CD28 for αβ T cells, and CD19, CD20, CD21, CD72 for mature B cells (Fig. 4a, Supplementary Table 1). Moreover, comparison between the above DEPs and DEGs shows that SPIDER detects canonical cell type markers which fail to be observed from RNA expression. For instance, CD21 (CR2) is a vital mature B cell marker, playing a role in antigen recognition and B cell activation (Ahearn, Joseph M., and Douglas T. Fearon., 1989; Fearon, Douglas T et al., 2000). In the HCC dataset, SPIDER’s prediction shows CD21 as a marker for mature B cells with distinct high expression at protein level, whereas RNA expression fails to unveil CR2 as a mature B cell marker due to its extremely low expression level (Fig. 4f).

Another example is CD16 (FCGR3A), an important NK cell marker initiating cellular toxicity (Lanier, Lewis L et al., 1988). SPIDER’s prediction successfully reveals CD16 as an NK-like cell marker, but this is not observed from RNA levels (Fig. 4b). These results prove that SPIDER can accurately predict the canonical cell type surface markers at protein level, including those that fail to be identified from RNA expression.

Besides the known canonical cell type markers, SPIDER also predicts some novel cell type markers which have not been reported by present studies, such as CD244 and CD94 for γδ1 T cells (Supplementary Table 1), suggesting SPIDER’s large potential in discovering novel markers that could be further used for defining and annotating various cell types.

One contribution of SPIDER to cell type annotation is discovering cell populations that could be overlooked by observing RNA expression alone. To prove this, we first cluster all cells based on highly confident surface proteins predicted by SPIDER, and then use canonical cell type markers from identified DEPs to annotate these clusters (Extended Data Fig. 6a). In comparison, we mask the pre-annotated cell type labels and re-cluster all cells based on their transcriptomes, and use canonical cell type markers from identified DEGs to annotate these clusters (Extended Data Fig. 6b, Methods). Our results show more subtly annotated cell subpopulations within the αβ T cell population with SPIDER’s prediction compared to transcriptomes, as a cluster of CD4^+^ CD25^-^ T cells is uniquely discovered with SPIDER’s prediction but not with transcriptomes (Extended Data Fig. 6c-e). Taken together, our results show that SPIDER assists with defining and annotating cell types.

### SPIDER facilitates identification of disease biomarkers on cell surface

Another important application of single-cell surface protein abundance is to unearth disease-related surface markers within a cell population, which could potentially serve as biomarkers for disease diagnosis and targets for disease treatment. To illustrate how SPIDER facilitates identification of disease-related surface markers, we use the same human liver scRNA-seq dataset with 2811 surface proteins predicted by SPIDER as previously described, and perform DE analysis for each cell population between tumor and control condition based on RNA levels or SPIDER-predicted protein levels (Supplementary Table 1).

Results of DEPs and DEGs reveal multiple HCC-related surface markers in immune cells, including those that could only be identified from SPIDER but not RNA expression. For instance, in mature B cells, CD40 is identified by SPIDER to be a positive surface marker for HCC, while RNA expression fails to show this (Fig. 4f). This prediction of SPIDER aligns with a recent publication, where CD40 is reported to be more highly expressed in tumor-infiltrating B cells compared to PBMC-derived B cells as measured by flow cytometry (Hladíková, Kamila et al., 2019). Other publications support the significance of SPIDER’s prediction by showing CD40’s vital role in regulating B cells and its great potential in HCC diagnosis and treatment (Sugimoto, Kazushi et al., 1999; Inoue, Satoshi et al., 2006). Other examples include SPIDER’s identification of PIK3IP1 as a negative, and PD-1 (CD279, PDCD1) as a positive surface marker within γδ2 T cells in HCC, a distinction not captured by RNA expression (Fig. 4g). SPIDER’s prediction of PD-1 is supported by previous publications where PD-1’s expression in γδ2 T cells is reported to be upregulated following antigenic stimulation in breast cancer as measured by flow cytometry (Iwasaki, Masashi et al., 2011). For PIK3IP1, although the relation between its expression in γδ2 T cells and HCC has not been reported yet, SPIDER’s prediction could be partially supported by the knowledge that T cells downregulate PIK3IP1 expression following activation (Saravia, Jordy et al., 2020), indicating that SPIDER facilitates the identification of potential novel disease markers such as PIK3IP1 which may be a novel negative HCC-related surface marker in γδ2 T cells.

By combining SPIDER’s prediction on cell type markers and disease-related markers together, we can even pinpoint disease biomarkers within more finely defined immune cell subpopulations. For instance, in αβ T cells, SPIDER identifies CD278 (ICOS) and CD177 as two positive surface markers for HCC (Fig. 4h). More specifically, SPIDER’s prediction clearly shows that CD278 is mainly highly expressed in the CD4^+^ T cell subpopulation in tumor environment, and CD177 is primarily expressed in the regulatory T cell (Treg) subpopulation as defined by canonical Treg markers CD4 and CD25. In contrast, CD177 is not observed to be an HCC-related marker from RNA expression (Fig. 4h). Recent publications strongly support SPIDER’s prediction by reporting higher CD278 expression in tumor-infiltrating CD4+ T cells (Di Blasi, Daniela et al., 2020), as well as upregulated CD177 expression in Tregs as measured via flow cytometry in HCC (Kim, Myung-Chul et al., 2021).

Besides immune cells, SPIDER can also identify disease-related surface markers in non-immune cells. For instance, in a hepatocyte subpopulation, SPIDER identifies CD44 as a positive HCC-related surface marker, which is not identified from corresponding RNA expression due to its extremely weak expression signal (Fig. 4c). SPIDER’s prediction aligns with previous publications reporting the upregulation of cell surface CD44 expression by malignant hepatocytes in HCC as tested via immunohistochemistry experiments (MATHEW, JOSEPH et al., 1996; Endo, Kanenori and Tadashi Terada., 2000).

Taken together, the results demonstrate that SPIDER facilitates the identification of disease-related cell surface markers, including not only those that have already been validated by previous publications, but also highly-potential novel markers that have not been reported such as PIK3IP1 (Fig. 4g). Moreover, SPIDER identifies disease-related surface markers that are overlooked by observing RNA expression alone, thereby opening up new avenues for novel disease biomarker discovery.

### Application to cell-cell interaction analysis

Cell-cell interactions (CCIs) mainly describe the physical interactions mediated by proteins and ligands between two or more cells. Analyzing CCIs helps elucidate communication networks and signaling pathways that govern cellular behavior, development, immune response, and disease progression, providing valuable insights into the heterogeneity and dynamics of cellular populations. Due to the lack of technology for proteomics measurement at single-cell resolution, existing single-cell CCI analysis tools such as CellChat only infers from transcriptomes based on ligand-receptor co-expression (Jin, Suoqin, et al., 2021; Armingol Erick, et al., 2021). With SPIDER-predicted single-cell surface protein abundance, we can now use CellChat to conveniently and directly infer CCIs from protein abundance data.

To showcase SPIDER’s application to CCI analysis, we use CellChat to infer CCIs from SPIDER-predicted highly confident cell surface proteins’ abundance on the previously described HCC dataset, and compare inferred CCIs between the normal and tumor environment (Methods). Our results show strengthened interactions between the mature B cell population (outgoing) and all the T cell populations (incoming) in tumor environment (Fig. 4i), supporting the notion that the functional interaction between T cells and B cells enhances local immune activation in HCC and contributes to better prognosis (Garnelo Marta, et al., 2017). Moreover, our results show a strengthened interaction between inflammatory macrophages (outgoing) and αβ T cells (incoming) in tumor environment (Fig. 4j), aligning with previous research indicating that inflammatory macrophages express T helper cell-attracting chemokines and promote T helper cell response upon antigen presentation (Biswas Subhra K and Alberto Mantovani, 2010).

Furthermore, our results unveil a weakened interaction between mature B cells (outgoing) and portal endothelial cells (incoming) in tumor environment, which has not been reported, suggesting that SPIDER may facilitate the discovery of novel cell-cell interactions in HCC (Fig. 4i).

Taken together, SPIDER facilitates the identification of CCIs, not only containing those that have already been validated by previous publications, but also potential novel CCIs that have not previously been reported. As a result, SPIDER may ultimately enhance our understanding of immune response and cell crosstalk in diseases.

### SPIDER facilitates the identification of cancer metastasis surface markers

To further showcase SPIDER’s broad application across various contexts, we delve into the study of cancer metastasis, a complex and dynamic process involving the spread of cancer cells from the primary tumor to distant organs or tissues. With the predicted surface protein abundance across multiple tissues by SPIDER, we aim to identify surface markers associated with cancer metastasis. To demonstrate this capability, we apply SPIDER to single-cell transcriptomes from a colorectal cancer (CRC) liver metastasis study containing a total of 125,150 cells (Che Li-Heng et al., 2021) (Fig. 5d-e). There are six datasets with distinct tissue and disease conditions (Extended Data Table 1). We use SPIDER to predict the abundance for as many as 2664 human cell surface proteins on these six transcriptome datasets (Supplementary Table 2).

**Fig. 5.**
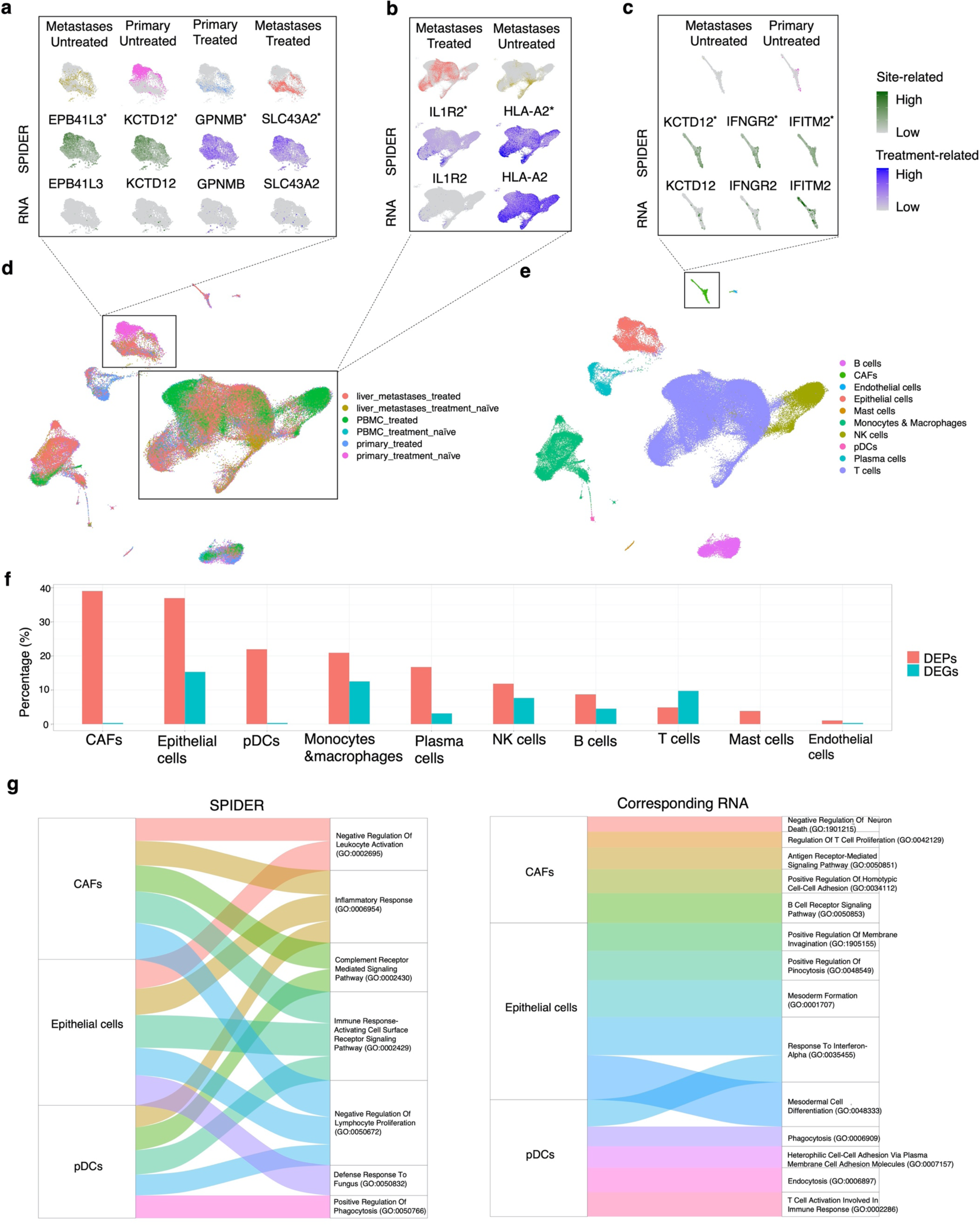
Application of SPIDER to human colorectal cancer liver metastasis scRNA-seq data. **a-e**, comparison between SPIDER-predicted surface protein abundance and corresponding RNA expression level for identified tissue site-related or treatment-related proteins within specific cell populations. Site-related surface proteins and their corresponding RNAs are colored dark-green, and treatment-related surface proteins are colored dark-purple. Surface proteins and genes labeled with asterisk mean they are DEPs or DEGs under corresponding differential expression analysis. **a-c**, UMAP visualization of predicted surface protein abundance and corresponding RNA expression for (**a**) epithelial cells, (**b**) T cells and (**c**) cancer-associated fibroblasts. **d**, UMAP visualization of all cells in the CRC liver metastases dataset, colored by contexts of tissue and treatment. **e**, UMAP visualization of all cells in the CRC liver metastases dataset, colored by cell types. **f,** comparison of the overall changes in SPIDER-predicted surface protein abundance to the overall changes in RNA expression during CRC metastasis within every cell population. For each cell population, DEPs and DEGs are identified between the CRC primary site and liver metastases based on the predicted protein abundance and RNA expression, respectively. The percentage of DEPs within a cell population is the number of identified DEPs divided by the number of the total predicted highly confident proteins; and from their corresponding genes, the percentage of DEGs within a cell population is also calculated. **g**, comparison between enriched pathways generated from SPIDER-predicted surface protein abundance and from corresponding transcriptomes in CAFs, epithelial cells, and pDCs.

To show that SPIDER enables the identification of metastasis-related surface markers, we take all treatment-naïve cells and conduct DE analysis for every cell population between different tumor sites based on RNA levels or SPIDER-predicted protein levels (Supplementary Table 2). Results of DEPs and DEGs reveal multiple metastasis-related surface markers in epithelial cells (cancer cells) that could only be identified from SPIDER’s prediction but not RNA expression. For instance, in epithelial cells, SPIDER identifies EPB41L3 and KCTD12, which are more lowly expressed in liver metastases site compared to the primary tumor site, as two negative surface markers for CRC liver metastasis; whereas they cannot be identified from the RNA expression (Fig. 5a). SPIDER’s prediction of EPB41L3 is supported by a study highlighting its downregulation in gastric cancer liver metastases, which significantly promotes cancer cell migration and invasion (Li Xiaohua et al., 2011). While direct evidence linking KCTD12 to cancer metastasis is currently absent, a recent study by Merhi Maysaloun et al partially supports our results, where decreased KCTD12 expression in CRC cells is reported to increase cancer cell viability and invasiveness (Merhi Maysaloun et al., 2023), indicating that KCTD12 could potentially be a novel CRC metastasis surface marker identified by SPIDER.

Besides cancer cells, SPIDER further identifies potential novel metastasis-related surface markers in other cell populations as well. For instance, in cancer-associated fibroblasts (CAFs), SPIDER detects IFITM2 which is more highly expressed in liver metastases compared to the primary site, as a positive surface marker, and KCTD12 and IFNGR2 as two negative surface markers for CRC liver metastasis; whereas these three markers cannot be identified from the RNA level (Fig. 5c). IFITM2 is verified by FACS to be expressed on human fibroblasts and contribute to cancer metastasis (De Marco, Margot, et al.). On the other hand, neither of the two predicted negative markers has been reported, suggesting SPIDER may have identified novel CRC metastasis-related surface markers.

We are also curious about the overall changes in surface protein abundance within every cell population during CRC liver metastasis, and how much difference there is between the overall changes in surface protein abundance and RNA expression. More significant changes in overall surface protein abundance may indicate larger contribution for that cell population to cancer metastasis. To explore this, we first summarize the total number of SPIDER-predicted highly confident proteins, from which we then calculate the proportion of DEPs between the primary tumor site and the liver metastases site for every cell population. In comparison, for these SPIDER-predicted highly confident proteins we select their corresponding genes, and perform the same calculation for DEG proportions to these genes for every cell population. As a result, we find that the overall change in surface protein abundance exceeds that of RNA expression (Fig. 5f). Specifically, CAFs (39.02%), epithelial cells (36.93%) and plasmacytoid dendritic cells (pDCs) (21.95%) exhibit the most significant alterations in surface protein expression patterns as predicted by SPIDER, aligning with the knowledge that CAFs play a vital role in modulating cancer metastasis through production of growth factors (Sahai Erik et al., 2020), as well as the fact that colorectal tumorigenesis results from progressive transformation of epithelial cells. pDCs are the most important professional antigen presenting cells (APCs) with the broadest range of antigen presentation (Kambayashi Taku, and Terri M Laufer, 2014; Mitchell Dana, Sreenivasulu Chintala, and Mahua Dey, 2018), therefore, it is also reasonable that pDCs show a great change in overall surface protein abundance as predicted by SPIDER. In contrast, RNA expression patterns merely change 0.35%, 15.33%, and 0.35% for CAFs, epithelial cells and pDCs, respectively, suggesting that SPIDER detects expression changes at protein level that fail to be revealed by corresponding RNA expression (Fig. 5f). There are cell populations showing consistent magnitude of expression changes at both protein level and RNA level as well, such as endothelial cells and mast cells with the smallest expression change (<5%) at both protein level and RNA level, indicating their minor role in CRC metastasis (Fig. 5f).

Taken together, these results show that SPIDER’s prediction of changes in surface protein abundance during cancer metastasis is biologically significant, and reveal evident difference in expression changes between surface protein level and RNA level, suggesting that SPIDER could complement the limitation of transcriptomes in metastasis marker identification.

To gain further insights into what pathways the above significantly changed surface proteins involve during cancer metastasis, we conduct pathway enrichment analysis based on the previously obtained DEPs and DEGs, respectively, for the top 3 cell populations CAFs, epithelial cells, and pDCs (Fig. 5g, Methods). Results show that enriched pathways generated from the predicted down-regulated surface proteins in metastases are largely related to immune response and shared by the three cell populations, suggesting major participation of these cell populations in cancer metastasis via regulating immune response by changing relevant surface proteins’ expression, and active interactions among these cell populations (Fig. 5g). While enriched pathways generated from RNA levels fail to show this pattern of collaborative immune response pathways shared by the three cell populations. Taken together, these results show that SPIDER compensate for the limitations in transcriptomes by accurately predicting cancer metastasis surface markers, and further reveal their biological roles within specific cellular contexts during metastasis, providing new insights for cancer metastasis study.

### SPIDER promotes the identification of disease treatment targets on cell surface

We have demonstrated SPIDER’s great potential in identifying disease biomarkers including cancer metastasis markers on cell surface. Furthermore, SPIDER also facilitates the identification of potential disease treatment targets. Using the same CRC dataset, we conduct DE analysis for every cell population between treatment-naïve and after-treatment conditions (Supplementary Table 2). Our results reveal that SPIDER predicts epithelial cells to express higher GPNMB in chemotherapy-treated compared to treatment-naïve liver metastases (Fig. 5a), indicating GPNMB’s involvement in drug responses, and its inhibitors could be potentially repurposed to enhance metastatic CRC treatment subsequent to chemotherapy. Our prediction aligns with a report from a phase 2 trial where GPNMB-targeted antibody is used for cancer treatment (Kopp, Lisa M., et al., 2019). Additionally, SPIDER predicts epithelial cells to express lower SLC43A2 at chemotherapy-treated CRC primary site compared to treatment-naïve site (Fig. 5a). Since it is reported that the high expression of SLC43A2 on tumor cells is linked with the impairment of CD8+ T cell’s function (Du Wan et al., 2022), SPIDER’s prediction on SLC43A2 is reasonable, indicating that inhibitors targeting SLC43A2 on cancer cells may serve as a novel CRC treatment. On the other hand, both GPNMB and SLC43A2 fail to be identified as DEGs from RNA expression. Taken together, these results show that SPIDER can assist with the identification of potential cell surface target for disease treatment.

Additionally, SPIDER predicts two immune response related surface proteins, IL1R2 and HLA-A2, to be down-regulated in liver metastases after chemotherapy, indicating an altered immune microenvironment after chemotherapy and potential treatment response (Fig. 5b).

Moreover, we compare CCIs inferred from SPIDER-predicted surface protein abundance or transcriptomes between treatment-naïve and after-treatment conditions on this CRC dataset.

Results show that SPIDER strengthens CCI signals and shows clearer changes in CCIs compared to using RNA (Extended Data Fig. 7). For instance, SPIDER exhibits an obvious weakened interaction between epithelial cells (outgoing) with other major immune cells (incoming) at the primary tumor site after chemotherapy treatment compared to before treatment (Extended Data Fig. 7a). While with CCIs inferred from RNA expression, these changes are not as obvious as the ones shown by SPIDER. Moreover, SPIDER exhibits weakened interactions between T cells (outgoing) and other cell populations (incoming) including epithelial cells and pDCs at the primary tumor site after treatment compared to before treatment (Extended Data Fig. 7b). On the other hand, CCIs inferred from RNA expression show no interaction between T cells and epithelial cells before and after chemotherapy, and the magnitude of interaction between T cells and pDCs stays unchanged (Extended Data Fig. 7b). These results indicate that SPIDER facilitates CCI analysis by magnifying the CCI signals, showing clearer changes in CCIs, and revealing CCI patterns that cannot be inferred from RNA expression data.

## Discussion

We propose SPIDER, a context-agnostic zero-shot deep ensemble model taking single-cell transcriptomes and *context factors* as input, that predicts abundance for a large scale of cell surface proteins on a query scRNA-seq dataset. Users can easily implement SPIDER by either using our pretrained model and saved weights for direct prediction on transcriptomes, or train their own SPIDER model using new reference CITE-seq datasets. There are three major distinctions of SPIDER. Firstly, it introduces zero-shot learning method through context-specific protein-protein similarity, which not only enables model prediction for unseen proteins, but also considers the contextual specificity of protein expression patterns (Fig. 1b, Extended Data Fig. 4). Confidence-estimation by SPIDER ensures a high overall accuracy for predictable unseen proteins (Fig. 3). Secondly, SPIDER is also unique in that it fully considers the effects of environmental context on surface protein abundance patterns by adding one-hot encoded *context factors* to input (Fig. 1b). The underlying biological meaning is that as the context changes, the process of protein translation from mRNA and transportation onto cell surface is regulated differently, which may ultimately lead to the change in cell surface protein abundance. SPIDER captures such influence of contexts on surface protein expression, enhancing its generalization across diverse contexts (Fig. 2c). Thirdly, SPIDER adopts a deep ensemble architecture, enabling one model to accommodate two different approaches to predicting seen and unseen proteins (Fig. 1b). The knowledge learned from DNN’s training is transferred to assist with the zero-shot prediction approach, making the model highly efficient. This architecture also enables SPIDER to include multiple CITE-seq datasets for training and train the union of all proteins, which maximizes data utilization compared to other models that either only train the intersection of all proteins in multiple training sets, or only include one single-cell dataset for training.

We have shown SPIDER’s generalization ability to a variety of contexts and its superior prediction performance in terms of both seen and unseen proteins compared to other baselines (Fig. 2 and Fig. 3). We have also showcased SPIDER’s broad applications including cell type annotation, disease biomarker identification, treatment target identification, and cell-cell interaction, proving that SPIDER compensate for the limitations in the transcriptome measurement (Fig. 4 and Fig. 5). Beyond these applications, SPIDER holds immense potential in many other tasks. For instance, coupling SPIDER’s prediction with drug discovery efforts may identify novel cell surface targets for personalized medicine based on an individual’s single-cell transcriptome profile. In addition, leveraging protein-protein similarity may enable the design of an optimal panel of surface proteins for CITE-seq experiments, essentially creating a novel reference set that ensures accurate prediction for the entire surfaceome.

SPIDER has its limitations. Firstly, it cannot predict for proteins whose corresponding RNA expression is not measured in the query dataset, indicating that in addition to transcriptome data, other types of information on such proteins may be needed for them to be predictable. Secondly, although SPIDER makes overall accurate prediction and outperforms other baselines, there are still a small number of proteins that could not be predicted well, which is also shown by other baselines (Fig. 2c). This is likely caused by the complexity of regulation in some proteins or the biological instability of some proteins’ abundance on cell surface, for instance, some surface proteins could be degraded rapidly after its transportation to cell surface, leading to very dynamic abundance at different time points.

One future direction involves augmenting the reference set with additional datasets, thereby furnishing SPIDER with more diverse contexts. Our current construction of SPIDER’s reference set comprises five tissues, four diseases and 17 cell types. Although this already covers a variety of contexts, there remains room for improving prediction performance, especially on contexts that extremely differ from the reference set. This is evident from our examination of the COVID- 19 BALF query dataset in figure 2c. With the continued advancement of CITE-seq technologies, it is likely that experiments will span a wider range of contexts, which could be incorporated into the reference set. Another avenue for future exploration would be extending the zero-shot ensemble architecture of SPIDER to other technologies of cell surface protein and transcriptome measurement such as REAP-seq and spatial transcriptomics. The current version has already included one ABseq dataset in the reference set (Extended Data Table 1), suggesting the plausibility of SPIDER to be applied to other technologies. Finally, we could utilize SPIDER to predict for intracellular protein abundance when relevant experimental data is available for training.

## Methods

### Dataset preprocessing

For all the RNA expression data from CITE-seq, ABseq and scRNA-seq (Extended Data Table 1), low-quality cells with <200 or >2500 genes detected are removed, cells with high percentages (>30%) of UMIs from mitochondrial genes are also removed. Low-quality genes with <5 counts are removed. The RNA expression data are then normalized with Seurat’s LogNormalize method. All the ADT data from CITE-seq and ABseq are normalized with Seurat’s centered log-ratio (CLR) method.

### Cell type annotation

The scArches-SCANVI embedding method requires annotated cell type labels for the reference set as input. For this input, we use SingleR and celldex packages in R to automatically annotate the cells in the reference set.

For SPIDER’s training, prediction, as well as internal and external validations, SingleR and celldex are used to give a consistent annotation of cell types for all the reference and query transcriptomes. Cell types with <100 cells across all training datasets are removed from SPIDER training. We set the parameter *ref* to *HumanPrimaryCellAtlasData()* for celldex.

For the two case study analysis after SPIDER’s prediction, cells in the HCC datasets are integrated in Seurat using the FindIntegrationAnchors and IntegrateData functions with the top 50 PCA dimensions. The re-clustering of the transcriptomes and the clustering of predicted surface protein abundance are performed by the FindClusters function in Seurat with a resolution of 0.5. The FindConservedMarkers function is used for identifying DEPs and DEGs as cell type markers for each cell cluster. The αβ T cell population is annotated using TRBC1, TRBC2, and CD3 (CD3D), and subpopulations are annotated using: Tregs (CD4^+^, CD25^+^ (IL2RA^+^)), double-negative T cells (CD3^+^, CD4^-^, CD8^-^ (CD8A^-^)), CD8^+^ T cells (CD3^+^, CD8^+^, CD4^-^), CD4^+^ CD25^-^ T cells (CD3^+^, CD4^+^, CD25^-^, CD8^-^). For the CRC transcriptome datasets, cells are integrated and clustered using the same method as described previously. Cell types are annotated using the markers as described in the original paper (Che, Li-Heng, et al, 2021).

### Transcriptomes embedding

We use scArches-SCANVI (version: 0.4.0) to embed all preprocessed RNA expression data for SPIDER training and prediction (Lotfollahi, Mohammad, et al., 2022). Top 1000 highly variable genes are selected from the training datasets and used for reference embedding. We save the reference embeddings for later training of the SPIDER model. For query datasets, the same 1000 genes as training sets are selected to generate query embedding. Genes that only exist in training datasets but not in query datasets are imputed with zeros. For both the reference and query embeddings, we set the parameter *n_latent* to 128 to obtain 128- dimensional feature embeddings (For other selections of feature dimensions, refer to Extended Data Fig. 1).

### The SPIDER model

#### Training sets

For normalized RNA expression data derived from *m* CITE-seq datasets, they are merged by shared genes and embedded into a single matrix *G*, where the number of rows is the sum of all cells, and the number of columns is 128. Provided that there are *k* proteins (*aka* seen proteins) in the union of normalized Antibody-Derived Tag (ADT) data from the same *m* CITE-seq datasets, SPIDER split the normalized ADT data and re-concatenated them into *k* vectors denoted as *P*_*i*_, where *P*_*i*_ ∈ ℝ^*ni*^, *and i* ∈ {1, 2, 3, …, *k*}. Here *P*_*i*_ consists of values of the *i*th protein’s cell surface abundance in all corresponding measured *n*_*i*_ cells across all *m* datasets. For any *P*_*i*_, SPIDER takes a *n*_*i*_ × 128 matrix *G*_*i*_ (*i* ∈ {1, 2, 3, …, *k*}) from *G*, where the *n*_*i*_ cells in *G*_*i*_ are the same *n*_*i*_ cells as in *P*_*i*_.

#### Deep neural networks

*k* individual DNNs are built and trained, denoted as *D*_*i*_ (*i* ∈ {1, 2, 3, …, *k*}), where *D*_*i*_ outputs the imputed cell surface abundance values for the *i*th protein. One-hot encoding of cell type, tissue and disease labels for the *i*th protein in all trained *n*_*i*_ cells are denoted as *c*_*i*_, *t*_*i*_, *d*_*i*_, respectively. During the training of *D*_*i*_, it learns a function *f*_*ij*_ mapping the protein abundance value in the *j*th cell *P*_*ij*_from [*G*_*ij*_ *c*_*ij*_ *t*_*ij*_ *d*_*ij*_], where the [] symbol denotes the row-wise concatenation operation. *f* denotes all such functions *fi* (*i* ∈ {1, 2, 3, …, *k*}) . *D*_*i*_’s hidden layers have the dimensions: 64, 32, 16. All layers are fully connected (FC) with rectified linear unit (ReLU) activation function (except the last layer). The objective function for *D*_*i*_ is

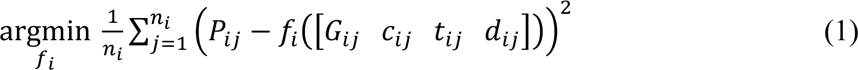

For each *D*_*i*_, the objective function is optimized stochastically with Adam with learning rate set to 0.0001 and internal 10-fold random holdout validation, where a random 10% of all *n*_*i*_cells are selected to be the internal validation set and the rest of the *n*_*i*_ cells are used as the training set.

For a given query dataset which is a *u* × *g* matrix denoted as *Q*, where *g* genes’ RNA expression values are measured in *u* cells, its embedded form is a *u* × 128 matrix denoted as *Q*′. One-hot encoding of cell type, tissue and disease labels in all *u* cells are denoted as *c*′, *t*′, *d*′, respectively. The surface abundance value of the *i*th seen protein in the *j*th cell is predicted as

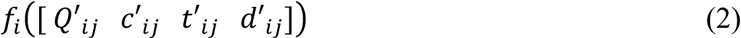

#### Protein-protein similarity

Given that there are *g*′ genes shared between the training transcriptomes and the query dataset *Q*, and provided there are *r* unseen proteins which we aim to predict for *Q* (To be predictable, these *r* unseen proteins should have their corresponding coding genes’ RNA expression measured in *Q*), and among all *k* seen proteins there are *k*′ proteins having their corresponding coding genes’ RNA expression measured in *Q*. For any *v*th seen protein (*v* ∈ {1, 2, 3, …, *k*′}), or any *t*th unseen protein (*t* ∈ {1, 2, 3, …, *r*}), the gene co-expression values are computed between its corresponding coding gene’s RNA expression (denoted as vector *Q*_(_ for any *v*th seen protein, or *Q*_)_ for any *t*th unseen protein) and every other genes’ RNA expression (denoted as *Q*_*w*_, *w* ∈ {1, 2, 3, …, *g*′}), where *Q*_(,_*Q*_),_*Q*_*_ ∈ ℝ^*u*^ and *u* is the aforementioned number of cells in *Q*. Denoting gene co-expression values as vector *Z*_*v*_ for any *v*th seen protein, or *Q*_t_ for any *t*th unseen protein, where *Q*_*v*_*Q*_*t*_*Q*_*w*_ ∈ ℝ^*u*^. The *w*th element in *Z*_*v*_ or *Z*_*t*_ is calculated as

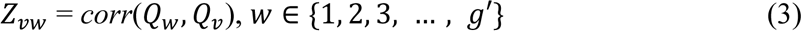

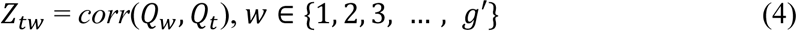

where *corr* denotes the computation of Pearson correlation.

Protein-protein similarity between the *v*th seen protein and the *t*th unseen protein is defined as the cosine similarity:

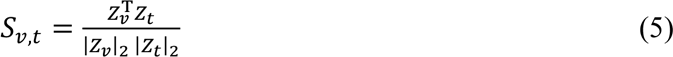

#### Ensemble member selection

A quality control is performed on DNNs via removing low-quality DNNs with internal validation performance of < 0.6 Pearson correlation coefficient between the prediction and ground truth. For any *t*th unseen protein, all DNNs are first ranked by their corresponding protein similarity (*S*) to this unseen protein, only the most relevant *N* DNNs with the largest protein similarity values are selected as ensemble members for predicting the *t*th unseen protein.

To decide the value of *N*, we design the following internal validation scheme: We consider every one of all *k* proteins in the reference set as an unseen protein. To create this setting, for the *i*th protein’s prediction (*i* ∈ 1,2,3, …, *k*), we set aside its corresponding trained DNN *D*_*i*_, and from the rest of all (*k*-1) DNNs, we rank their corresponding proteins’ similarity (*S*) to the *i*th protein from the largest to the smallest. We then select the top *N* DNNs with the largest *S* values to form ensemble members for predicting the *i*th protein. In this case, the set-aside *i*th protein is an unseen protein whose data will not be used for its prediction. We consider the set-aside *i*th protein’s paired transcriptome and protein abundance data used for training *D*_*i*_ as an internal validation set, and use the selected *N* ensemble members to make prediction on the set-aside protein’s transcriptome data as described in *Zero-shot deep ensembles*. For every one of all *k* proteins, we repeat this set-aside and prediction procedure, and compare its predicted protein abundance to the actual protein abundance to obtain a Pearson correlation coefficient, then we calculate the average correlation across all *k* proteins. For any *N* varying from 1 to *k*, we repeat the above steps, and plot the average correlation across all *k* proteins. We find that the internal validation performance is the best when *N*=8 (Extended Data Fig. 8), therefore, we set *N* to 8 for SPIDER.

#### Zero-shot deep ensembles

For the *t*th unseen protein, the ensemble model linearly combines all the separate predictions made by the *N* ensemble members, respectively. Each ensemble member’s corresponding protein similarity (*S*) serves as its linear coefficient. *S*_2,)_ denotes the value of protein similarity between the *l*th ensemble member’s corresponding (seen) protein and the *t*th unseen protein. A softmax-like normalization is performed to constrain *S* to non-negative values:

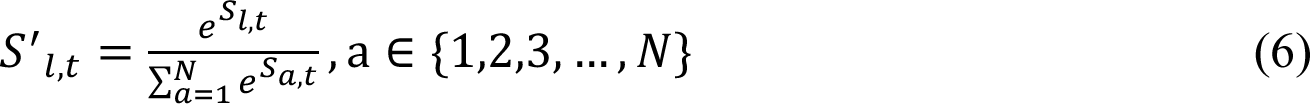

The output from the ensemble model is the final predicted value for the unseen protein’s cell surface abundance. The surface abundance value of the *t*th unseen protein in the *j*′^th^ cell in the query dataset *Q* is predicted as

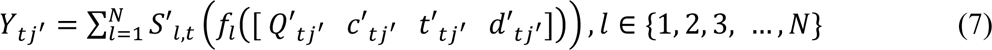

where *f*_*i*_ denotes the mapping function learned from the *l*th ensemble member out of all *N* ensemble members.

#### Prediction confidence estimation

The *t*th unseen protein is estimated to obtain highly confident prediction if

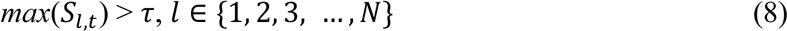

We set τ as 0.85 in our study. A seen protein is estimated to obtain highly confident prediction if it has DNN internal validation performance of > 0.6 Pearson correlation coefficient between the prediction and ground truth.

### Benchmarking procedures

Seurat anchor V4 imputes surface protein abundance via finding a set of reference-based transfer anchors between the query and the reference datasets (Hao, Yuhan, et al., 2021). totalVI imputes surface protein abundance via learning protein likelihood parameters from a variational autoencoder (VAE) that encodes cells into a latent space (Gayoso, Adam, et al., 2021). For multiple CITE-seq datasets as a combined reference set, Seurat V4 and totalVI originally only support imputation prediction of the intersection of all proteins for a new query transcriptome dataset (i.e., by default, proteins which do not overlap between multiple CITE-seq datasets cannot be trained, and cannot be predicted either). sciPENN is a deep learning model enabling multiple CITE-seq datasets which may have incompletely overlapped proteins to serve as reference. By default, all the proteins in union can be trained via sciPENN’s censored loss function approach, and can be imputed for a new query transcriptome dataset. To ensure a fair comparison with SPIDER in terms of predicting for seen proteins, we use the same preprocessed reference data as SPIDER for Seurat V4, totalVI and sciPENN, where there are *m* CITE-seq datasets with *k* proteins in union, and merge the *m* RNA expression matrices by shared genes (*sg*) without further embedding. We also use the same four external validation sets as SPIDER (Extended Data Table 1) for model evaluation. For Seurat’s and totalVI’s training, we first follow the same method as described in **The SPIDER model**’s *Training datasets* to split the transcriptome data and the normalized protein abundance data, obtaining split transcriptome matrix *G*′_*i*_ (*i* ∈ 1,2,3, …, *k*) of dimensions *n*_*i*_ × *sg* and protein abundance vectors *P*_*i*_ (where *P*_*i*_ ∈ ℝ^*ni*^, *i* ∈ {1, 2, 3, …, *k*}) for the *i*th protein. Then Seurat V4 and totalVI are implemented to impute the *i*th protein for the query transcriptome dataset. We perform the above implementation one time for imputing one seen protein for one query dataset until finishing the imputation of all the seen proteins for all four query datasets. For sciPENN’s training, we directly input the reference data without splitting them.

cTPnet is a pretrained multi-task DNN model for imputing cell surface abundance for 24 proteins, with no option for training on new data (Zhou, Zilu, et al., 2020). For cTP-net, we use its saved weights to directly predict on the same query datasets as SPIDER. We only compare the shared predicted seen proteins between SPIDER and cTP-net. The training set for cTPnet is different from SPIDER’s reference dataset. Therefore, this comparison is not a direct head-to-head between SPIDER and cTPnet. Also, since the healthy bone marrow dataset among our query datasets was used for cTPnet’s pretraining, we exclude this query dataset from benchmarking comparison with cTPnet.

Pearson correlation and root mean square error (RMSE) are used as metrics to evaluate the prediction accuracy between the predicted abundance and its ground truth pre-measured by CITE-seq. The metrics for SPIDER, Seurat and totalVI are all calculated in the log-normalized feature space.

### Gene ontology terms

GO terms are generated using R package ontologySimilarity for each tested unseen protein’s corresponding gene, and one-hot encoded. We select GO terms that appear at least one time across all tested unseen proteins. The one-hot encoded selected GO terms serve as protein representations to further derive protein-protein similarity.

### Protein-protein interaction score

Human PPI scores (with the default threshold of >400) are obtained from STRINGdb (Szklarczyk, Damian, et al., 2023). For each training protein (i.e., seen protein) or tested unseen protein, the PPI scores between that protein and all the other proteins listed in STRINGdb serve as the protein representation to further derive protein-protein similarity. Missing PPI scores are imputed with zeros.

### Downstream analysis using predicted surface protein abundance

Of all human cell surface proteins listed by the UniProt database, proteins present in SPIDER’s reference set are imputed by SPIDER as seen proteins. Among the rest of the proteins listed, proteins with their corresponding genes present in the query scRNA-seq dataset are imputed by SPIDER as unseen proteins. The SPIDER-imputed seen and unseen proteins together constitute the large-scale surfaceomes and these predicted abundance data are used for downstream analysis.

### Differential expression analysis

For identification of DEGs and DEPs in disease biomarker analysis and metastasis marker analysis, we use the FindMarkers function in Seurat, where the *logfc.threshold* parameter is set to 0.25, and the *test.use* parameter is set to “wilcox”. For identification of DEGs and DEPs in cell type annotation with the HCC and CRC datasets, we use the FindConservedMarkers function in Seurat. Genes and proteins with adjusted p values < 0.05 are considered DEGs and DEPs. DoHeatmap and FeaturePlot functions in Seurat are used for visualizing the proteins and genes.

### Cell-cell interaction analysis

The CellChat package in R is used to for cell-cell interaction analysis. For cell surface protein-based analysis, we replace the RNA expression data with SPIDER-predicted surface protein abundance data and repeat the same analysis procedures as using RNA expression data. As CellChat does not support negative values as input, we set all negative values in protein abundance data to zeros. For the identifyOverExpressedGenes function, we set the parameters *thresh.pc, thresh.fc, thresh.p* to 0.1, 0.25 and 0.05, respectively.

### Pathway enrichment analysis

Pathway enrichment analysis is performed by Seurat’s DEenrichRPlot function. We set the parameters *num.pathway, logfc.threshold, p.val.cutoff* to 5, 0.25, 0.05, respectively. The obtained *p* values are used to generate alluvial plot by ggplot2 in R.

### Statistical analysis

Wilcoxon rank-sum test (Mann–Whitney U-test) in Seurat is used for all DE analysis. *p* < 0.05 is considered significant.

## Data availability

Public datasets for training and prediction of SPIDER in this manuscript can be found at National Center for Biotechnology Information Gene Expression Omnibus (GEO) under accession number GSE164378 (GSM5008737, GSM5008738), GSE163120 (GSM4972212), GSE172155 (GSM5242790, GSM5242791, GSM5242792, GSM5242793), GSE143363, GSE128639, GSE165045 (GSM5025052, GSM5025059), GSE167118 (GSM5093918), GSE149614, GSE178318, as well as at figshare: https://figshare.com/projects/Single-cell_proteo-genomic_reference_maps_of_the_human_hematopoietic_system/94469, respectively.

## Code availability

The code for SPIDER is available at https://github.com/Bin-Chen-Lab/SPIDER.

## Author Contributions

B.C. conceived the study and supervised the study. R.C. performed the analysis. J.Z. contributed to machine learning method development. All authors contributed to the writing of the manuscript.

## Acknowledgements

We thank Dr. Rama Shankar for providing liver cancer single cell annotation data and the Chen lab members for the discussion. The research is supported by the NIH R01GM134307 and R01GM145700. The content is solely the responsibility of the authors and does not necessarily represent the official views of sponsors.

