## supplemental materials for "Imputing abundance of over 2500 surface proteins from single-cell transcriptomes with context-agnostic zero-shot deep ensembles"

| tissue | Disease state | # of cells trained | technology | # of proteins (Quantified or predicted) | # of patient | Use |
|---|---|---|---|---|---|---|
| Peripheral blood<br><br>(Hao, Yuhan, et al., 2021) | healthy | 89473 | CITE-seq | 228 | 8 | Training |
| Bone marrow<br><br>(Triana, Sergio, et al., 2021) | healthy | 9823 | ABseq | 105 | 1 | Training |
| Brain<br><br>(Pombo Antunes, Ana Rita, et al., 2021) | grade IV glioblastoma | 12360 | CITE-seq | 268 | 1 | Training |
| Pleura<br><br>(Ma, Xiaojun, et al., 2021) | malignant pleural mesothelioma | 2761 | CITE-seq | 52 | 1 | Training |
| Peritoneum<br><br>(Ma, Xiaojun, et al., 2021) | malignant peritoneal mesothelioma | 3941 | CITE-seq | 48 | 1 | Training |
| Bone marrow<br><br>(Pei, Shanshan, et al., 2020) | acute myeloid leukemia | 2103 | CITE-seq | 19 | 1 | Training |
| Bone marrow<br><br>(Stuart, Tim, et al., 2019) | healthy | 32588 | CITE-seq | 25 | 1 | External validation |
| Pancreas | pancreatitis | 1412 | CITE-seq | 13 | 1 | External validation |

| | | | | | | |
|---|---|---|---|---|---|---|
| (Lee, Bomi, et al., 2022) | | | | | | |
| Pancreas<br><br>(Lee, Bomi, et al.,2022) | healthy | 1239 | CITE-seq | 13 | 1 | External validation |
| Bronchoalveolar lavage fluid (BALF)<br><br>(Zhao, Yu, et al., 2021) | COVID-19 | 7387 | CITE-seq | 39 | 1 | External validation |
| Liver | hepatocellular carcinoma | 16709 | scRNA-seq | 2811 | 10 | Case study |
| Liver | healthy | 22211 | scRNA-seq | 2811 | 8 | Case study |
| Colon | Colorectal cancer, treatment-naïve | 28029 | scRNA-seq | 2664 | 3 | Case study |
| Liver | Colorectal cancer liver metastasis, treatment-naïve | 23448 | scRNA-seq | 2494 | 3 | Case study |
| Peripheral blood | Colorectal cancer, treatment-naïve | 12342 | scRNA-seq | 1991 | 1 | Case study |
| Colon | Colorectal cancer, treated | 19546 | scRNA-seq | 2612 | 3 | Case study |
| Liver | Colorectal cancer liver metastasis, treated | 27373 | scRNA-seq | 2597 | 3 | Case study |
| Peripheral blood | Colorectal cancer, treated | 14412 | scRNA-seq | 2038 | 2 | Case study |

**Extended Data Table 1. Summary of all data sets used.**

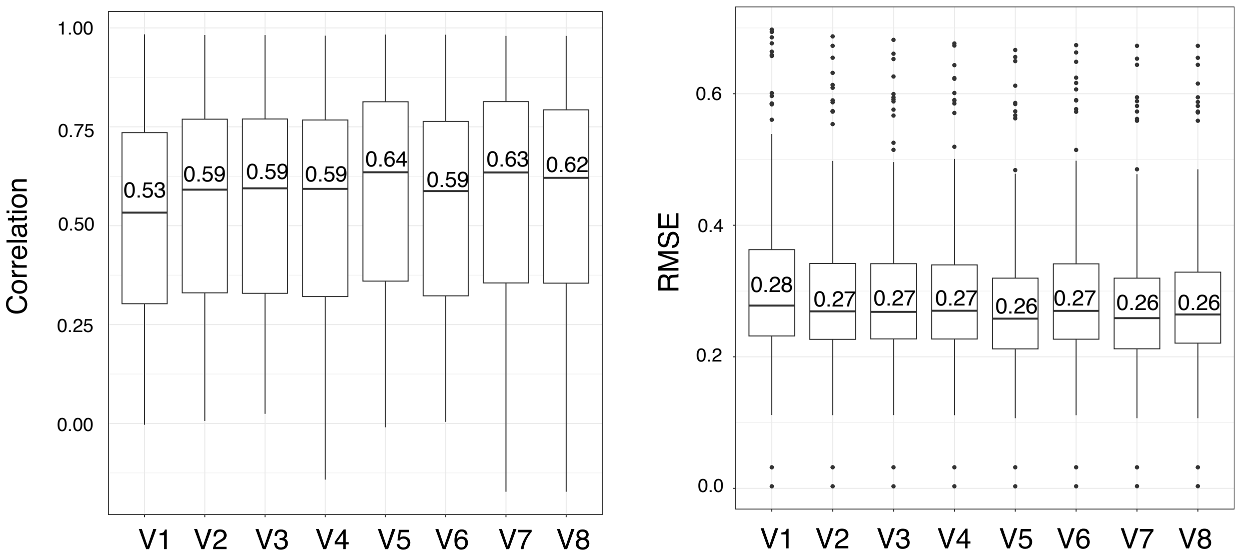


**Extended Data Fig. 1 | SPIDER DNN's Internal 10-fold random holdout validation performance for different model designs.** Evaluation metrics are Pearson correlations and RMSEs between each protein's prediction and ground truth across all cells. Median correlations and median RMSEs are compared among all model versions. V1: Only using 10-dimensional transcriptome embeddings as input for DNNs. V2: Only using 64-dimensional transcriptome embeddings as input for DNNs. V3: Only using 256-dimensional transcriptome embeddings as input for DNNs. V4: Only using 128-dimensional transcriptome embeddings as input for DNNs. V5: Using 128-dimensional transcriptome embeddings + Encoded information on tissue, disease and cell type as input for DNNs. V6: Using 128-dimensional transcriptome embeddings + Encoded information on cell type as input for DNNs. V7: Using 128-dimensional transcriptome embeddings + Encoded information on tissue as input for DNNs. V8: Using 128-dimensional transcriptome embeddings + Encoded information on disease as input for DNNs. V5 is used in SPIDER.

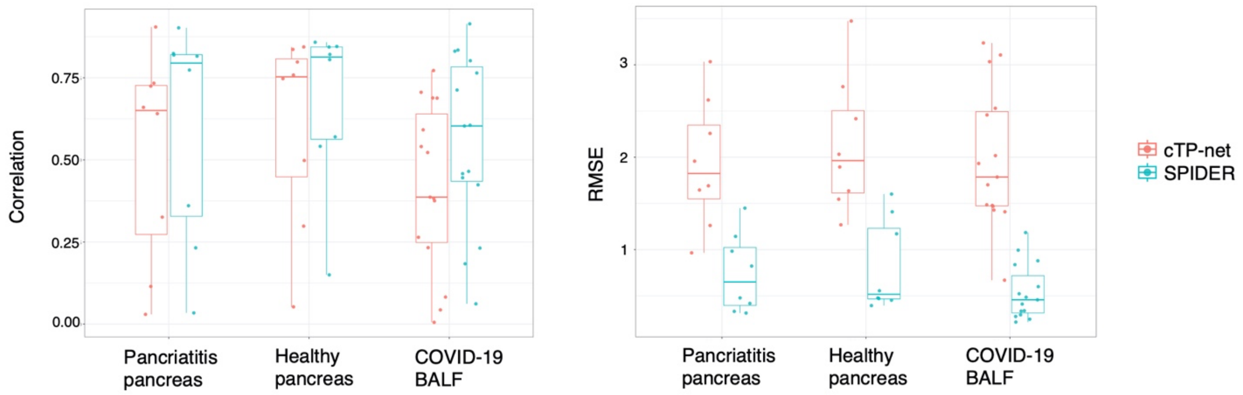


**Extended Data Fig. 2 | Comparison of external validation performance between SPIDER and cTP-net for predicting seen proteins.** Evaluation metrics are Pearson correlations and RMSEs between each protein's prediction and ground truth across all cells.

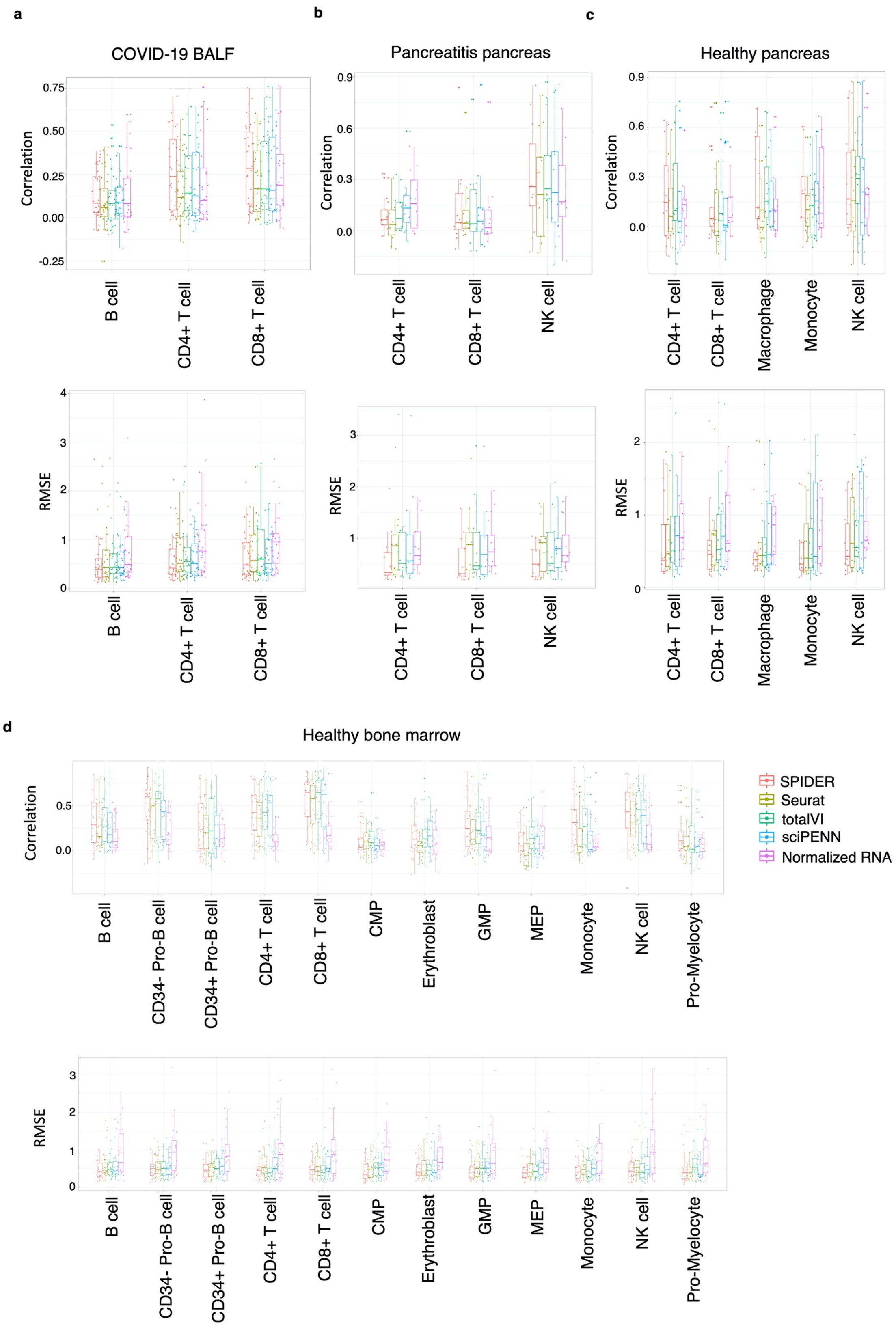


**Extended Data Fig. 3 | Benchmarking of within cell type prediction accuracy for every seen protein in four external validation datasets.** Cell types containing at least 100 cells are selected for evaluation. Pearson correlation and RMSE are shown between every cell surface protein's predicted abundance (or normalized RNA expression) and CITE-seq measured abundance for each method. Each dot represents a protein. **a**, Healthy bone marrow. **b**, pancreatitis pancreas. **c**, healthy pancreas. **d**, COVID-19 BALF.

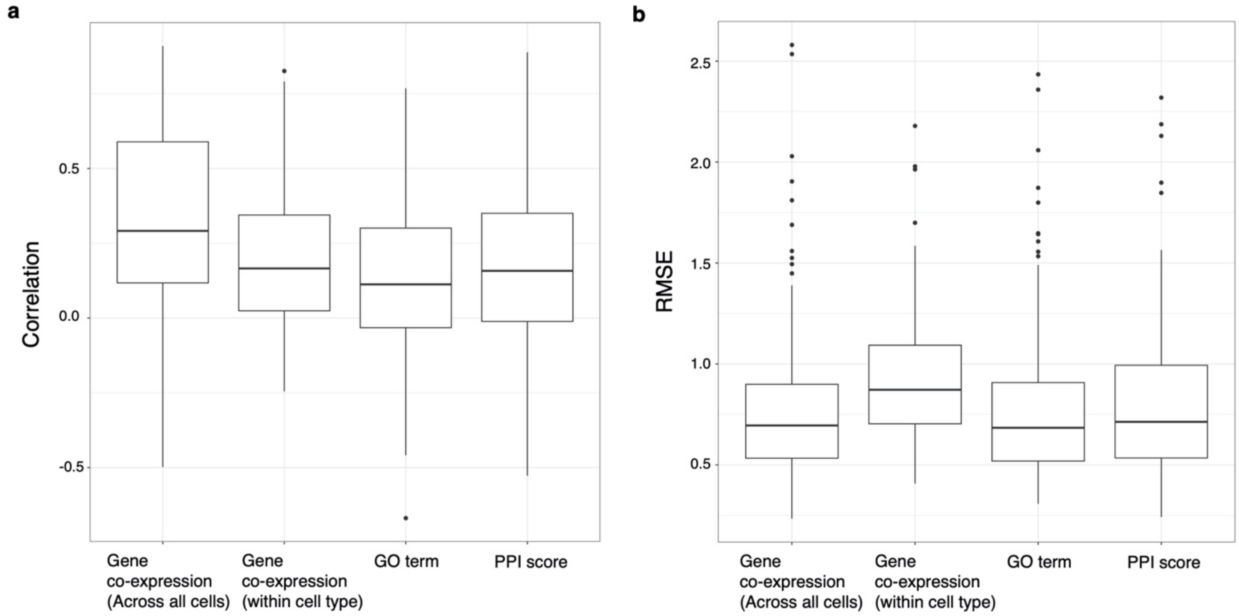


**Extended Data Fig. 4 | Comparison of model performance for unseen proteins' prediction using different types of protein representations for generating protein-protein similarity.** Internal validation performance for unseen proteins is evaluated by Pearson correlation and RMSE between each protein's predicted abundance and ground truth across all cells. **a**, using Pearson correlation. **b**, using RMSE.

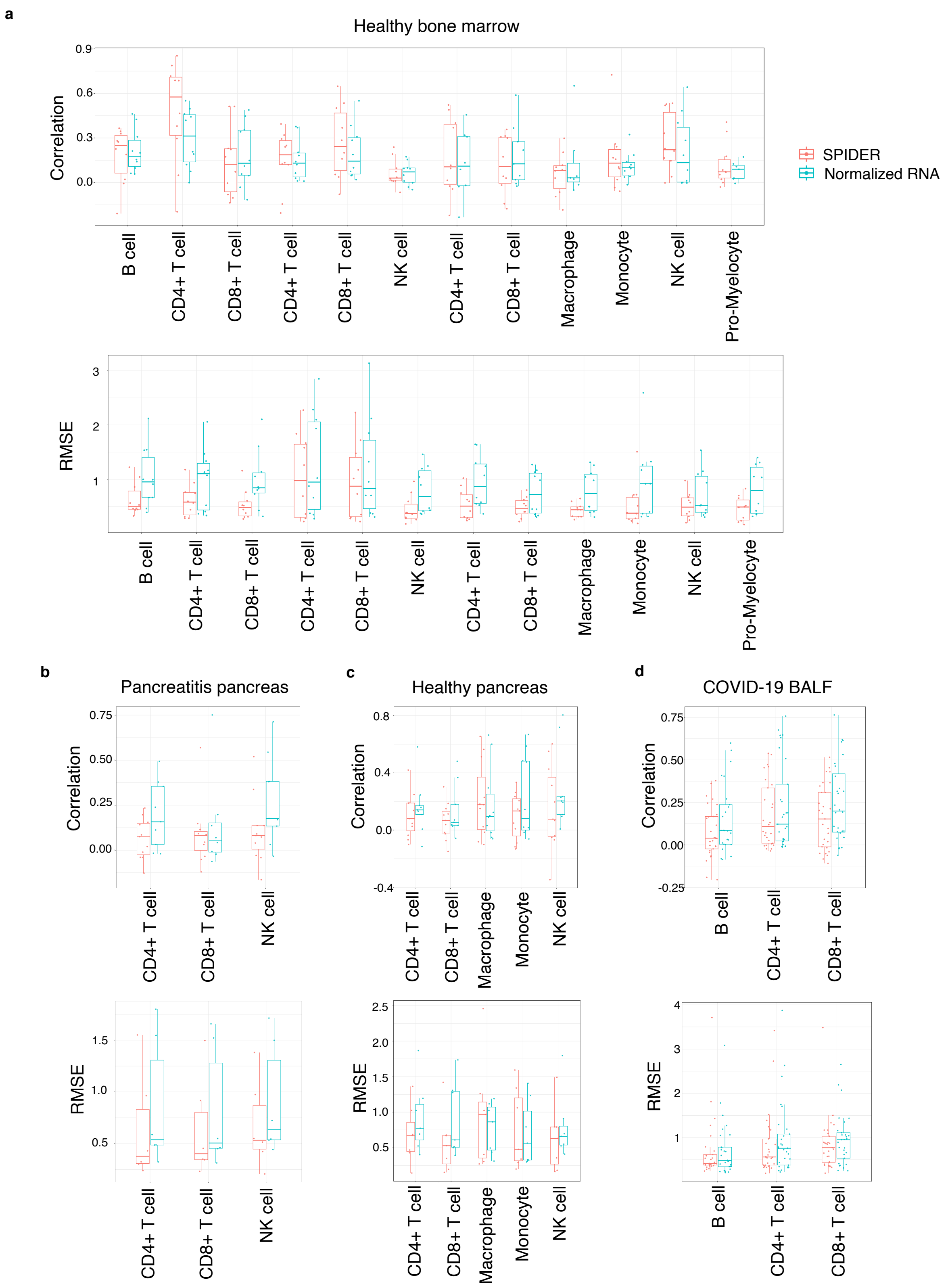


**Extended Data Fig. 5 | Benchmarking of within cell type prediction accuracy for every unseen protein in four external validation datasets.** Cell types containing at least 100 cells were selected for evaluation. Pearson correlation and RMSE are shown between every cell surface protein's predicted abundance (or normalized RNA expression) and CITE-seq measured abundance for each method. Each dot represents a protein. **a**, Healthy bone marrow. **b**, pancreatitis pancreas. **c**, healthy pancreas. **d**, COVID-19 BALF.

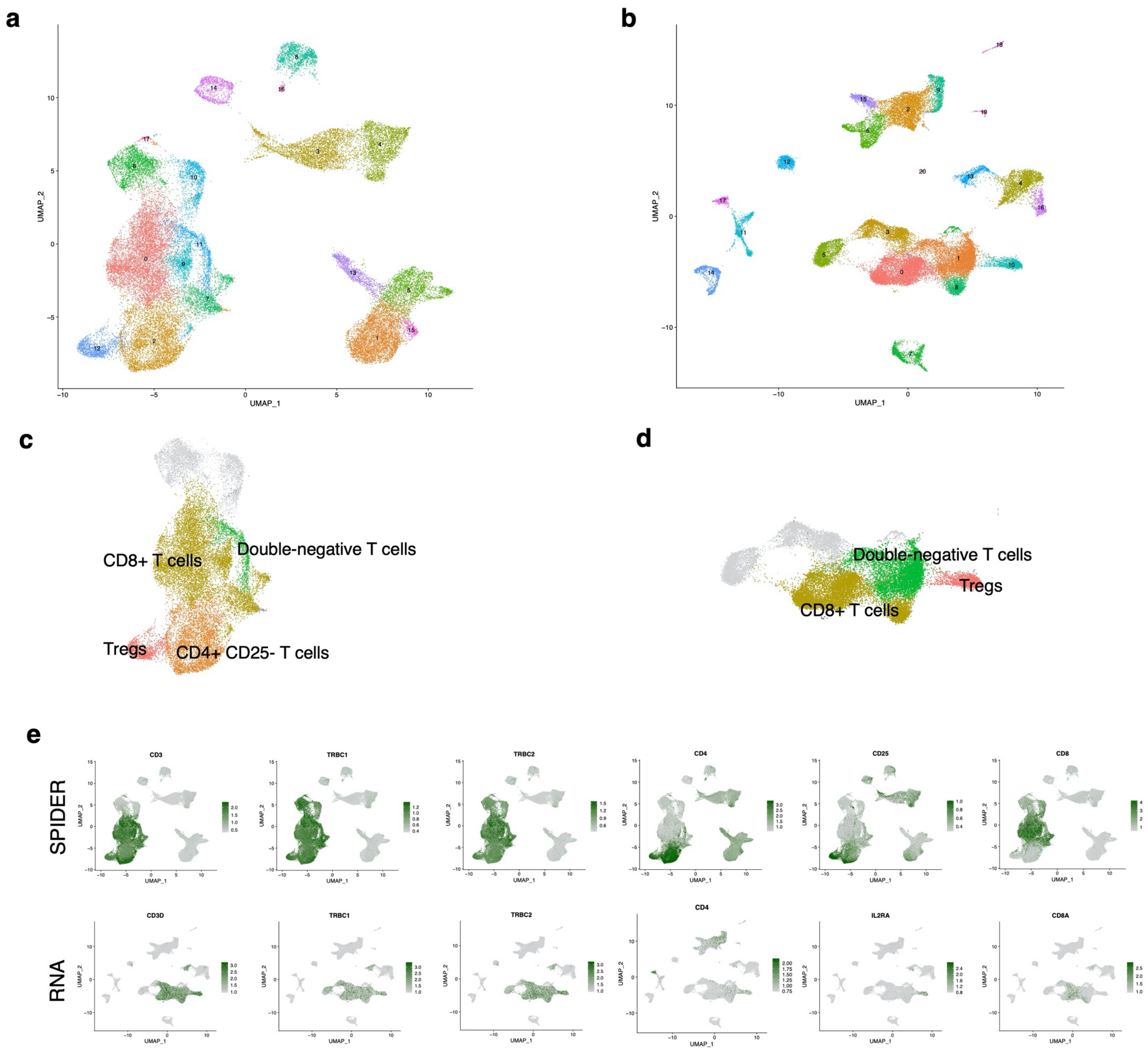


**Extended Data Fig. 6 | Comparison for cell type annotations between using transcriptome data or using SPIDER-predicted surface protein abundance alone in the HCC dataset. a,** UMAP visualization of all cell clusters in the HCC dataset using SPIDER-predicted surface protein abundance data. Only the abundance of estimated highly confident proteins is used for cell clustering. **b,** UMAP visualization of all re-clustered cells in the HCC dataset using transcriptome data. **c**, annotated subpopulations within the αβ T cell population using SPIDER-predicted surface protein abundance. **d**, annotated subpopulations within the αβ T cell population using transcriptome data. **e**, UMAP visualization of the SPIDER-predicted surface protein abundance and transcriptome data, colored by predicted protein abundance and normalized RNA expression of cell type markers, respectively.

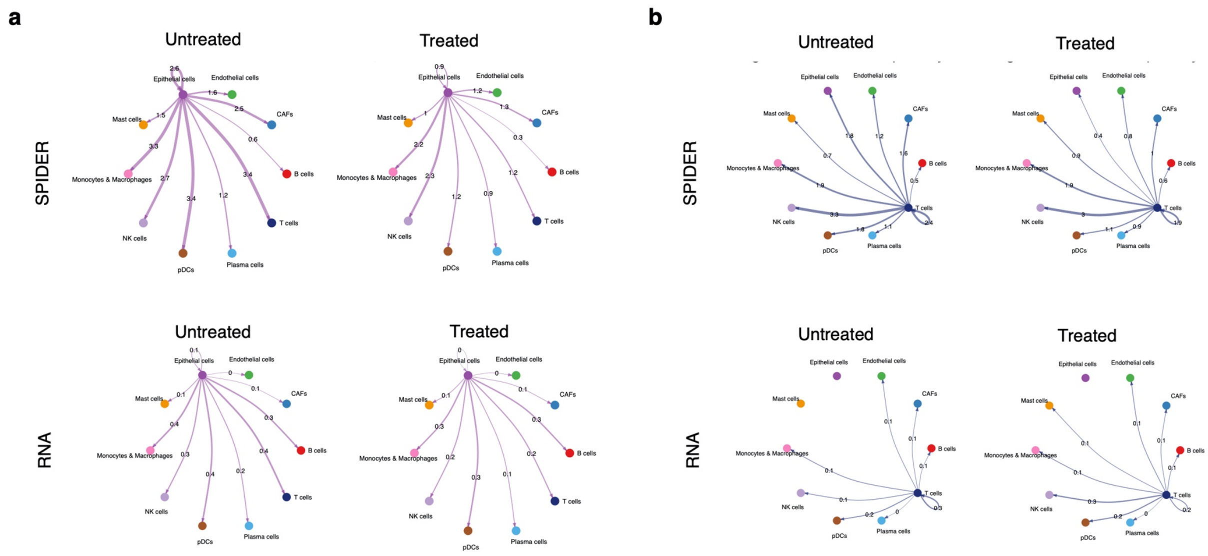


**Extended Data Fig. 7 | Comparison of CCIs inferred from SPIDER-predicted surface protein abundance to CCIs inferred from CRC liver metastases transcriptome data.** The values represent the strength of interactions. **a**, CCIs inferred from SPIDER-predicted surface protein abundance and transcriptomes between epithelial cells and other cell populations at the CRC primary site under treatment-naive and treated conditions. **b**, CCIs inferred from predicted surface protein abundance and transcriptomes between T cells and other cell populations at the CRC primary site under treatment-naive and treated conditions.

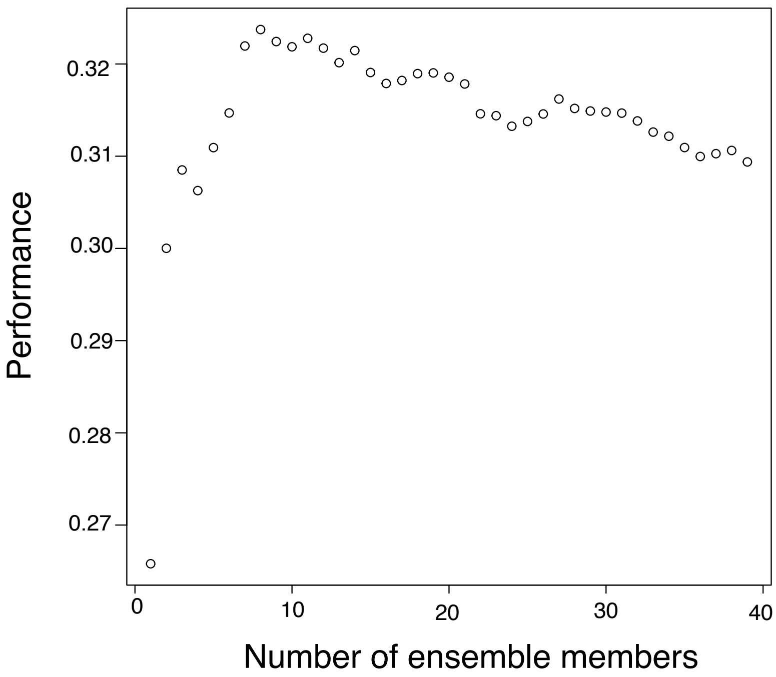


**Extended Data Fig. 8 | Internal validation performance of unseen proteins using different numbers of top ensemble members.** The internal validation performance is evaluated by the average Pearson correlation between every unseen protein's prediction and ground truth abundance.
